# Brain DNA methylomic analysis of frontotemporal lobar degeneration reveals *OTUD4* in shared dysregulated signatures across pathological subtypes

**DOI:** 10.1101/2022.10.21.513088

**Authors:** Katherine Fodder, Megha Murthy, Patrizia Rizzu, Christina E. Toomey, Rahat Hasan, Jack Humphrey, Towfique Raj, Katie Lunnon, Jonathan Mill, Peter Heutink, Tammaryn Lashley, Conceição Bettencourt

**Affiliations:** Queen Square Brain Bank for Neurological Disorders, UCL Queen Square Institute of Neurology, London, UK; Department of Neurodegenerative Disease, UCL Queen Square Institute of Neurology, London, UK; Department of Clinical and Movement Neurosciences, UCL Queen Square Institute of Neurology, London, UK; German Center for Neurodegenerative Diseases (DZNE), Tübingen, Germany; The Francis Crick Institute, London, UK; Nash Family Department of Neuroscience & Friedman Brain Institute, Icahn School of Medicine at Mount Sinai, New York, NY, USA; Department of Clinical and Biomedical Sciences, Faculty of Health and Life Sciences, University of Exeter, Exeter, UK; Alector, Inc., South San Francisco, CA, USA

**Keywords:** DNA methylation, frontotemporal dementia, progressive supranuclear palsy, human brain tissue, EWAS, co-methylation

## Abstract

Frontotemporal lobar degeneration (FTLD) is an umbrella term describing the neuropathology of a clinically, genetically and pathologically heterogeneous group of diseases, including frontotemporal dementia (FTD) and progressive supranuclear palsy (PSP). Among the major FTLD pathological subgroups, FTLD with TDP-43 positive inclusions (FTLD-TDP) and FTLD with tau positive inclusions (FTLD-tau) are the most common, representing about 90% of the cases. Although alterations in DNA methylation have been consistently associated with neurodegenerative diseases, including Alzheimer’s disease and Parkinson’s disease, little is known for FTLD and its heterogeneous subgroups and subtypes. The main goal of this study was to investigate DNA methylation variation in FTLD-TDP and FTLD-tau. We used frontal cortex genome-wide DNA methylation profiles from three FTLD cohorts (234 individuals), generated using the Illumina 450K or EPIC microarray. We performed epigenome-wide association studies (EWAS) for each cohort followed by meta-analysis to identify shared differential methylated loci across FTLD subgroups/subtypes. Additionally, we used weighted gene correlation network analysis to identify co-methylation signatures associated with FTLD and other disease-related traits. Wherever possible, we also incorporated relevant gene/protein expression data. After accounting for a conservative Bonferroni multiple testing correction, the EWAS meta-analysis revealed two differentially methylated loci in FTLD, one annotated to OTUD4 (5’UTR-shore) and the other to NFATC1 (gene body-island). Of these loci, OTUD4 showed consistent upregulation of mRNA and protein expression in FTLD. Additionally, in the three independent co-methylation networks, *OTUD4*-containing modules were enriched for EWAS meta-analysis top loci and were strongly associated with the FTLD status. These co-methylation modules were enriched for genes implicated in the ubiquitin system, RNA/stress granule formation and glutamatergic synaptic signalling. Altogether, our findings identified novel FTLD-associated loci, and support a role for DNA methylation as a mechanism involved in the dysregulation of biological processes relevant to FTLD, highlighting novel potential avenues for therapeutic development.

## INTRODUCTION

Frontotemporal lobar degeneration (FTLD) is an umbrella term describing the neuropathology of a group of neurodegenerative disorders, which are characterised by the selective degeneration of the frontal and temporal lobes of the brain. These disorders are clinically, pathologically and genetically heterogeneous. Clinically, patients with FTLD frequently present with frontotemporal dementia (FTD), which is the second most common form of early onset dementia and is often associated with behavioural and language changes. A fraction of patients may present with or develop Parkinsonism as part of their disease, including those with progressive supranuclear palsy (PSP), and frontotemporal dementia and parkinsonism linked to chromosome 17 (FTDP-17). An overlap with amyotrophic lateral sclerosis/motor neuron disease (ALS/MND) is also observed in a proportion of patients with FTLD, highlighting a spectrum of clinical phenotypes that relate to shared neuropathologic features [17, 41].

A considerable number of FTLD cases report a positive family history (30-50%), and the majority of familial cases can be attributed to mutations in three genes, namely chromosome 9 open reading frame 72 (*C9orf72*), progranulin (*GRN*), and microtubule-associated protein tau (*MAPT*). Apart from those cases in which a genetic mutation has been identified, neuropathological assessment is essential to confirm the disease entity underlying FTLD. The neuropathological classification of FTLDs, based on the presence/absence of specific proteinaceous inclusions, recognizes five major subgroups. FTLD with 43 kDa transactive response DNA-binding protein (TDP-43) positive inclusions (FTLD-TDP), and with tau positive inclusions (FTLD-tau), account for the vast majority of cases, representing around 50% and 40% of FTLD cases, respectively [27, 58].

Even though progress has been made in identifying genetic risk factors for diseases under the FTLD umbrella [19, 23, 34, 66, 87], the molecular mechanisms driving FTLD pathology are not completely understood. Mounting evidence reveals changes in the FTLD brain transcriptional landscapes [3, 15, 28, 30, 81]. However, studies investigating non-sequence-based regulatory mechanisms such as epigenetic modifications in FTLD brain tissue are limited [9, 51, 83, 86]. Variable DNA methylation, the most well studied epigenetic modification, has consistently been associated with Alzheimer’s disease pathology in epigenome-wide studies (EWAS) and subsequent meta-analyses [72, 74, 89]. In FTLD, brain tissue EWAS are scarce and limited to a single PSP study [83].

To investigate further the relevance of DNA methylation variation in FTLD, we set out a study investigating epigenome-wide DNA methylation variation in frontal lobe tissue from three cohorts, spanning different subtypes of FTLD-TDP and FTLD-tau subgroups, followed by an EWAS meta-analysis, co-methylation network analysis in each cohort, and subsequent module preservation analysis in the other datasets. Through the EWAS meta-analysis we identified two differentially methylated loci shared across the FTLD subgroups and subtypes after a conservative Bonferroni correction for multiple testing. These methylation sites were annotated to *OTUD4* (5’UTR-shore) and *NFATC1* (gene body-island). We also identified co-methylation modules associated with the FTLD status, FTLD subtypes, and pathological features (e.g., brain atrophy and severity of neuronal loss). Functional and cellular enrichment analyses have shown an overrepresentation of gene ontology terms related to regulation of gene expression and the ubiquitin system as well as specific cell types, including pyramidal neurons and endothelial cells, across FTLD subgroups and subtypes. In all three independent co-methylation networks, OTUD4-containing modules were enriched for top EWAS meta-analysis loci, and were strongly associated with the disease status, further supporting their role in FTLD. Our findings implicate DNA methylation in the dysregulation of important processes in FTLD, including the ubiquitin system, RNA/stress granule formation and glutamatergic synaptic signalling.

## METHODS

### Demographic and clinical characteristics of post-mortem brain donors

For FTLD cohort 1 (FTLD1, N=23), all post-mortem tissues originated from brains donated to the Queen Square Brain Bank archives, where tissues are stored under a licence from the Human Tissue authority (No. 12198). Both the brain donation programme and protocols have received ethical approval for donation and research by the NRES Committee London – Central. All cases were characterized by age, gender, disease history (including disease onset and duration) as well as neuropathological findings. For FTLD cohort 2 (FTLD2, N=48), all post-mortem tissues were obtained under a Material Transfer Agreement from the Netherlands Brain Bank, and MRC Kings College London, as described by Menden et al. [53]. For FTLD cohort 3 (FTLD3, N=163, after quality control), data made available by Weber et al. [83] was retrieved from GEO (accession code GSE75704). Figure 1 shows an outline of the study design and analysis framework. More details on each cohort are presented in Table 1.

**Table 1:**
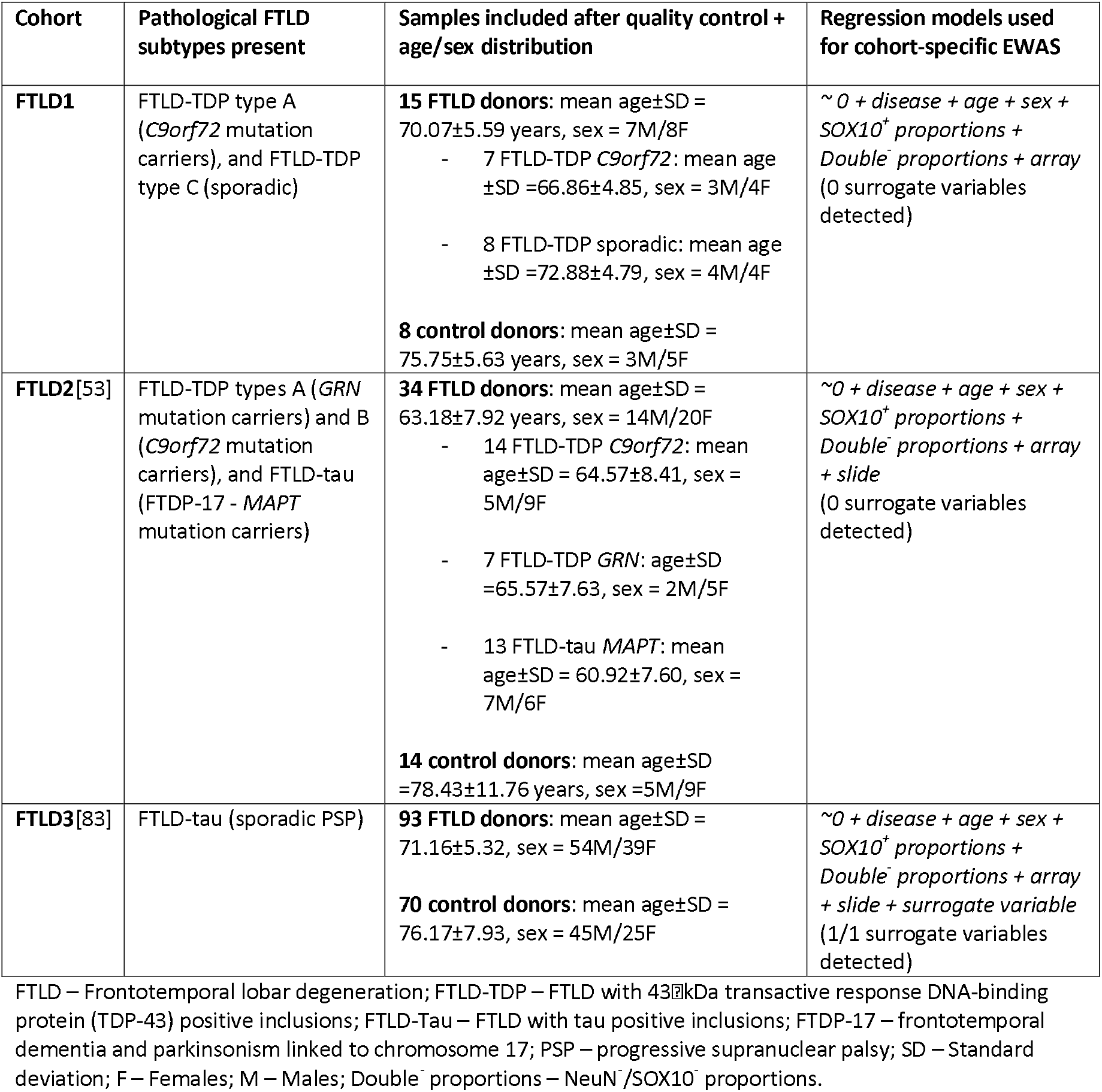
Clinical and pathological characteristics of the three FTLD cohorts and selected models for cohort-specific EWAS.

**Figure 1.**
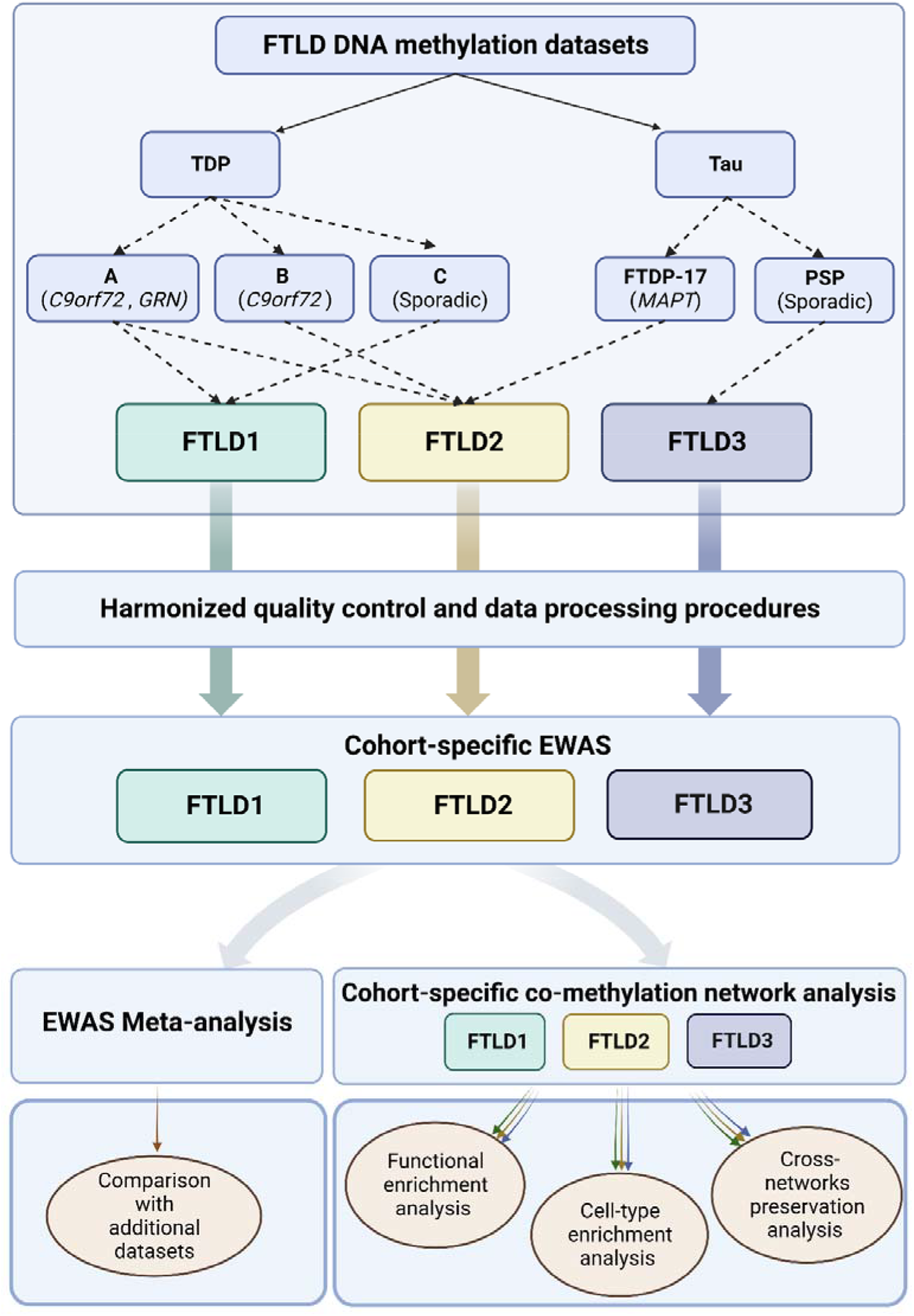
Outline of the study design and analysis framework. FTLD – Frontotemporal lobar degeneration; PSP – Progressive supranuclear palsy. Figure created with BioRender.

### Measures of brain atrophy, neuronal cell loss and pathology related traits

For FTLD1 and a proportion of cohort FTLD2, formalin-fixed paraffin-embedded (FFPE) sections were also available for more detailed neuropathological evaluations, including sections stained for standard haematoxylin and eosin (H&E). These FFPE sections were from the opposite brain hemisphere with respect to the frozen tissue used for the DNA methylation profiling.

For FTLD1 and FTLD2, microscopic atrophy was assessed on H&E stained slides, by examining the cortical thickness and neuronal loss in the frontal and temporal cortices. A four point grading system was used in comparison to a neurological normal control with no underlying neurodegenerative changes: 0 – the cortical thickness was within normal limits and no neuronal loss was observed; 1 – reduction in cortical thickness but the number of neurons was comparable to normal levels; 2 – reduction in cortical thickness and reduction in the numbers of neurons; 3 – severe reduction in cortical thickness and no neurons observed. For each region, the microscopic atrophy was scored semi-quantitatively by an experienced observer blinded to clinical, histopathological and genetic status, at an objective magnification of×20. Macroscopic atrophy was also determined for FTLD1 based on observations of gyri and sulci from the coronal slices observed during brain cutting procedures. Levels of atrophy were graded, as previously described [69], into four stages: none, mild, moderate, and severe. These neuropathological scores of the frontal and temporal regions were used in the module-trait correlations with the DNA co-methylation network modules.

### DNA methylation profiling and data quality control

For FTLD1, genomic DNA was extracted from carefully dissected flash frozen frontal cortex grey matter tissue using standard protocols. Bisulfite conversion was performed with the EZ DNA Methylation Kit (Zymo Research) using 500 ng of genomic DNA. For FTLD2 and FTLD3, DNA extractions and bisulfite conversions were performed previously as described by Menden et al. [53] and Weber et al. [83]. Genome-wide methylation profiles were generated using the Infinium HumanMethylationEPIC BeadChip (Illumina) for FTLD1 and FTLD2, or the Infinium HumanMethylation450 BeadChip (Illumina) for FTLD3, as per the manufacturer’s instructions.

Beta values ranging from 0 to 1 (approximately 0% to 100% methylation, respectively), were used to estimate the methylation levels of each CpG site using the ratio of intensities between methylated and unmethylated alleles. Data analysis was conducted using several R Bioconductor packages as previously described [11]. All three cohorts were subjected to harmonized quality control checks and pre-processing. Briefly, raw data (idat files) were imported and subjected to rigorous pre-processing and thorough quality control checks using minfi [4], wateRmelon [67], and ChAMP packages [76]. The following criteria were used to exclude probes that did not pass quality control checks from further analysis: 1) poor quality, 2) cross reactive, 3) included common genetic variants, and 4) mapped to X or Y chromosome. In addition, samples were dropped during quality control if: 1) they presented with a high failure rate (≥ 2% of probes), 2) the predicted sex did not match the phenotypic sex, and 3) they clustered inappropriately on multidimensional scaling analysis. Beta values were normalised with ChAMP using the Beta-Mixture Quantile (BMIQ) normalisation method. M-values, computed as the logit transformation of beta values, were used for all statistical analysis, as recommended by Du et al. [22], owing to their reduced heteroscedasticity (as opposed to beta-values) and improved statistical validity for differential methylation analysis.

As significant batch effects were detected during quality control checks, and different FTLD subgroups/subtypes were studied in FTLD1-3, the three cohorts were analysed separately first and then meta-analysed. Similarly, co-methylation network analyses were conducted on each cohort separately, and module preservations were then cross-checked with data from the other cohorts (as described in more detail below).

### Cell type deconvolution based on DNA methylation data

As DNA methylation patterns are often cell-type specific, changes in different brain cell-type proportions constitute an important confounding factor for DNA methylation studies performed on ‘bulk’ brain tissue. We used a novel cell-type deconvolution reference panel recently described by Shireby et al. [72] which brings more granularity and expands previous methods that account only for neuronal (NeuN+) versus all other cell types (NeuN-). This new method uses novel DNA methylation data obtained from fluorescence activated sorted nuclei from cortical brain tissue to estimate the relative proportions of neurons (NeuN+), oligodendrocytes (SOX10+) and other glial brain cell types (Double-[NeuN-/SOX10-]). Cell-type proportions in bulk brain tissue were thus estimated using the CETYGO (CEll TYpe deconvolution GOodness) package (https://github.com/ds420/CETYGO), and the sorted cell-type reference datasets as described by Shireby et al. [72]. Pairwise comparisons between FTLD cases and controls were conducted using Wilcoxon rank sum test with Benjamini-Hochberg correction for multiple testing, and adjusted p<0.05 was considered significant.

### Differential methylation analysis and EWAS meta-analysis

We applied linear regression models (Table 1) using the M-values as the input to identify associations between DNA methylation variation at specific CpG sites and FTLD using the limma package [64]. For FTLD1, we have accounted for possible confounding factors, such as age and sex as well as factors detected in principal components 1 and 2 as seen in Singular Value Decomposition (SVD) plots (ChAMP package), which included cell proportions (SOX10+ and Double-) and sample position in the array. Using this regression model, no surrogate variables were detected with the num.sv function of the SVA package [42], meaning there were no remaining unknown, unmodelled, or latent sources of noise [64]. The same process was applied to FTLD2 and FTLD3. The model for FTLD2 was further adjusted for slide, whereas for FTLD3, the model was further adjusted for slide and one surrogate variable (Table 1). False discovery rate (FDR) adjusted p-values <0.05 were considered genome-wide significant.

We used the estimated coefficients and SEs obtained from the regression models, described above for the three FTLD cohorts, to undertake an inverse variance meta-analysis using the metagen function from the meta R package [8]. Only methylation probes present in all datasets (N= 363,781) were considered for this analysis. When reporting differentially methylated sites, a conservative Bonferroni significance was defined as p<1.374×10^−7^ (p < 0.05/363,781) to account for multiple testing. We report random-effects meta-analysis results as the three cohorts included different FTLD subgroups/subtypes according to the neuropathological classification possibly leading to high heterogeneity in the meta-analysis. We also used a less stringent FDR adjusted p < 0.10 to report top meta-analysis loci, all of which were then investigated in the co-methylation networks.

### Co-methylation network analysis

To identify clusters of highly correlated CpGs (co-methylation modules) in an unsupervised manner, i.e., agnostic of gene ontology, we used a systems biology approach based on weighted gene correlation network analysis (WGCNA) [38]. For this analysis, we focused on CpGs present in all three FTLD datasets, non-intergenic CpGs (i.e. CpGs annotated to genes), and selected the top 20% with the highest variance across individuals in each cohort regardless of their disease status (i.e., most variable 56,001 CpG sites per cohort). After outlier exclusion, a total of 23, 42 and 157 samples remained in the FTLD1, FTLD2 and FTLD3 cohorts, respectively. For each network, we used as input the M-values adjusted for the covariates included in the models described above (Table 1) and constructed signed networks. Modules were calculated using the WGCNA blockwiseModules function, with a minimum module size of 200 and a soft-thresholding power of 16, 10 and 12 for the FTLD1, FTLD2 and FTLD3 networks, respectively. Module membership (MM) was then reassigned for each network using the applyKMeans function of the CoExpNets package [13]. Highly connected CpGs within a module (hub CpGs) present with high MM values to the respective module. In the results section, we refer to hub CpGs as those with the highest MM within a given module.

By using a principal component analysis on the CpG methylation values within each module, the CpGs inside each module were represented by a weighted average, the module eigengene (ME). The MEs were then correlated with the FTLD status, FTLD subtypes, and other sample traits, including disease onset and duration, measures of macroscopic atrophy and neuronal loss scores, and other pathology related traits, as available for each cohort.

To gain insights into the biology underlying the FTLD-related modules, we carried out functional enrichment for CpGs mapping to genes using the default parameters of clusterProfiler [85]. We also carried out cell-type enrichment analysis on the FTLD-related modules using the package EWCE [73] and associated single-cell transcriptomic data [88].

### DNA methylation cross-network module preservation analysis

As a method for differential network analysis, i.e., to identify which co-methylation modules in each of the three generated FTLD networks were preserved (i.e., shared) or perturbed (i.e., unique) in the other two datasets, we employed module preservation analysis, as described by Langfelder et al. [39] . For each network (taken as the “reference dataset”), module preservation in the other two datasets (the “test data”) was calculated using the modulePreservation function implemented in WGCNA. In all instances, the “test data” contained methylation values (adjusted M-values) for the 56,001 CpG sites used to construct the “reference dataset” network. A total of 200 permutations for each preservation analysis was used. As a measurement of module preservation, we used the Z-summary statistic (a composite measure to summarise multiple preservation statistics). A Z-summary greater than 10 indicates a strong preservation of this module in the “test data”, a Z-summary of between 2 and 10 indicates moderate preservation, and a Z-summary less than 2 indicates no preservation.

### Comparisons of DNA methylation hits with FTLD frontal/temporal cortex gene expression data

To examine the gene expression patterns of the EWAS meta-analysis gene hits, we used previously published transcriptomics data from bulk frontal cortex tissue of FTLD-TDP cases and controls [30] as well as bulk temporal cortex tissue of FTLD-tau cases (PSP) and controls [82]. To further infer the expression patterns of selected DNA methylation hit genes in specific brain cell types, we also correlated gene expression levels (adjusted for age, sex, and RNA integrity number) with cellular proportions using data from Hasan et al. [30], with cellular proportions estimated using the method described by Mathys et al. [50]

### Comparisons of DNA methylation hits with FTLD-TDP frontal cortex proteomics data

To examine the gene expression patterns of the EWAS meta-analysis gene hits at the protein level, we used proteomics data from FTLD-TDP and controls. Briefly, frontal cortex homogenate of frozen post-mortem human brain tissue was prepared from control (N=6), FTLD-TDP type A with *C9orf72* repeat expansion (N=6), and FTLD-TDP type C (N=6) cases. Proteins in both the soluble supernatant and the insoluble pellet fraction were analysed, and samples were pooled per disease group (three cases per pooled sample). Proteins were quantitated using 2D-LCMS and UDMSe label-free proteomics and SYNAPT G2-Si High Definition mass spectrometer operating in ion mobility mode. Data was processed using Progenesis software, as previously described [78]. A total of 6114 proteins were detected in the supernatant, and 5108 in the pellet, with an overlap in some proteins that were found both in the supernatant and pellet. Fold-changes between FTLD-TDP subtypes compared to controls were calculated. Of the Bonferroni significant EWAS meta-analysis hits, only the OTUD4 protein was detected (both in the supernatant and in the pellet).

### Comparisons of DNA methylation hits with additional datasets

We further investigated the normal expression patterns of the meta-analysis gene hits both in the human and mouse brains using single nuclei RNAseq data from the Allen Brain Map (https://celltypes.brain-map.org/)[7], and data from the Allen Mouse Brain Atlas (http://mouse.brain-map.org) [43]. Given the *OTUD4*-related findings, we investigated the list of cortical tissue OTUD4 protein interactors made available by Das et al. [20]. The RNA granule database (http://rnagranuledb.lunenfeld.ca/) collates curated literature evidence that support gene or protein association with the stress granules (SGs) and P-bodies (PBs). We used a list of tier 1 genes from the RNA granule database version 2.0 for comparisons with the lists of genes composing the three *OTUD4* FTLD-associated co-methylation modules.

### OTUD4 immunohistochemical staining

To investigate tissue expression patterns of OTUD4 protein across the human cortex, FFPE frontal cortex tissue from 7 FTLD cases (4 FTLD-TDP type A and 3 FTLD-TDP type C) and 3 controls (overlapping with FTLD1) were utilized. Briefly, eight-micrometre-thick sections cut from the FFPE blocks were immunostained using a standard avidin-biotin-peroxidase complex method with di-aminobenzidine as the chromogen [40]. The rabbit anti-OTUD4 antibody (Atlas Antibodies HPA036623, 1:200) was used, along with heat antigen retrieval pre-treatment prior to application of the primary antibody. The samples were mounted and examined using a light microscope.

## RESULTS

### Cell-type deconvolution based on DNA methylation data highlights important cellular composition differences in FTLD

To estimate brain cell-type proportions in our bulk frontal cortex DNA methylation datasets, we used a refined cell-type deconvolution algorithm based on reference DNA methylation profiles from purified nuclei from neurons (NeuN+), oligodendrocytes (SOX10+) and other brain cell types (NeuN-/SOX10-)[72]. This new model controls better for cellular heterogeneity in bulk cortex tissue compared to previous models, which account only for neuronal (NeuN+) versus all glial cells (NeuN-). Within each sample group, we observed extensive variability in cell-type proportions across cell types (Fig. 2). When comparing disease cases with controls, no overall differences were overall detected in the proportions of oligodendrocytes (SOX10+) and other glial cells (NeuN-/SOX10-) after accounting for multiple testing corrections. However, with the exception of the PSP cases (FTLD3), all FTLD subgroups/subtypes showed a significant decrease in neuronal proportions compared to controls (Wilcoxon rank sum test, adjusted-p<0.05), as expected in neurodegenerative diseases. These findings highlight the importance of adjusting for cell-type proportions in bulk tissue EWAS studies. Accounting for this allowed us to identify DNA methylation changes that are relevant to the disease rather than merely reflecting changes in cell-type composition, which could be related partly to the disease pathogenesis itself and partly due to technical issues (e.g., a result of capturing different proportions of grey and white matter during tissue dissection).

**Figure 2.**
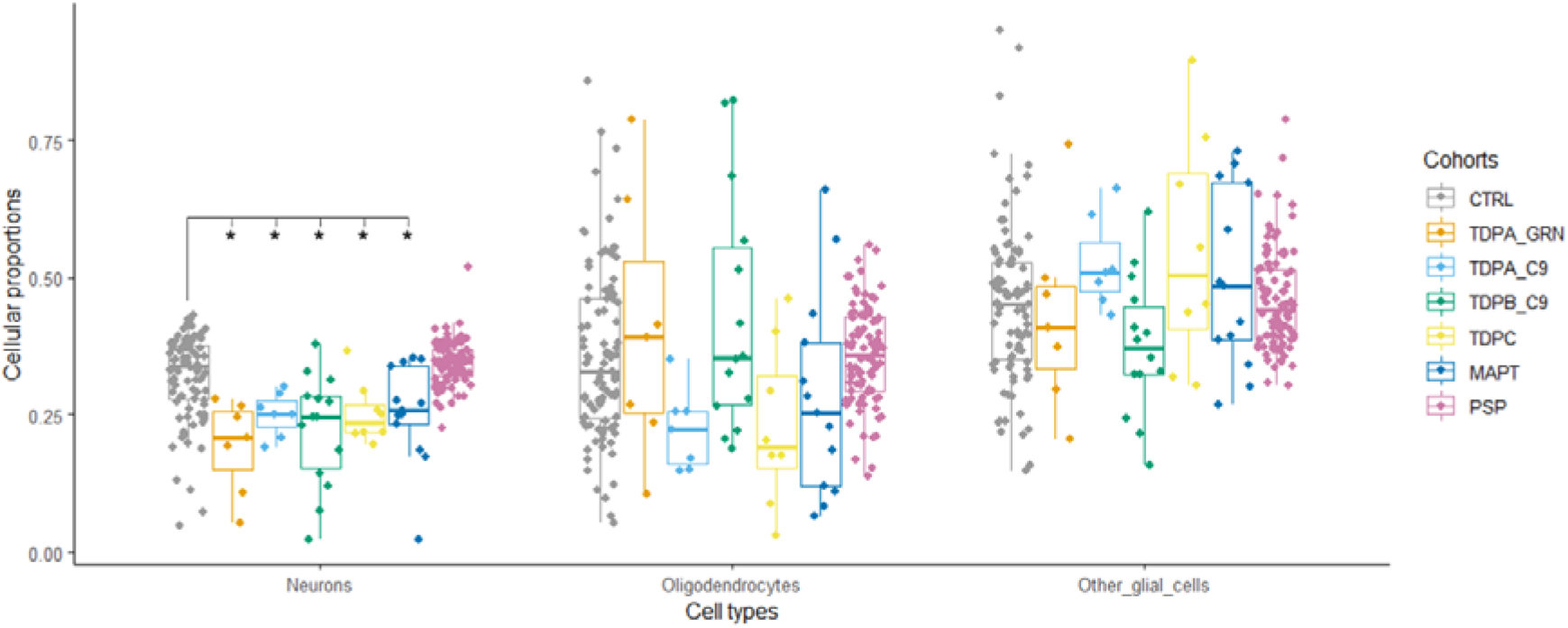
Brain cell-type proportion estimates derived from bulk DNA methylation data in frontal lobe of frontotemporal lobar degeneration (FTLD) and controls. *indicates significant differences for each cell-type between FTLD subtypes and the corresponding controls; pairwise comparisons were performed using the Wilcoxon rank sum test, and adjusted p-values <0.05 we considered significant. CTRL – controls; TDPA_GRN – FTLD with TDP-43 positive inclusions (FTLD-TDP) subtype A, carriers of *GRN* mutations; TDPA_C9 – FTLD-TDP subtype A, carriers of *C9orf72* repeat expansion; TDPB_C9 – FTLD-TDP subtype B, carriers of C9orf72 repeat expansion; TDPC – FTLD-TDP subtype C, sporadic; *MAPT* – FTLD with tau positive inclusions (FTLD-Tau), carriers of MAPT mutations; PSP – FTLD-Tau, sporadic progressive supranuclear palsy (PSP); Neurons – NeuN+; Oligodendrocytes – SOX10+; other glial cells – NeuN-/SOX10-.

### Frontal cortex case-control EWAS meta-analysis identifies shared differentially methylated CpG sites across FTLD pathological subgroups and subtypes

First, we investigated DNA methylation variation in specific loci across the genome as covered by the 450K/EPIC arrays, using linear regressions models to perform cohort-specific case-control EWAS. For FTLD1 and FTLD2, which comprise heterogeneous cases with sporadic and genetic forms of FTLD-TDP and FTLD-tau pathology, no genome-wide significant CpGs were identified. For FTLD3, which only includes cases with FTLD-tau pathology (sporadic PSP), 234 differentially methylated positions were identified (Supplementary Table S1, Online Resource). The top differentially methylated CpG in the FTLD3 cohort was cg09202319, which was hypomethylated in FTLD-tau (PSP) compared to controls (adjusted-p=6.54 x 10^-8^). This CpG mapped to a CpG island in the promoter region of PFDN6 (Prefoldin Subunit 6), which is involved in promoting the assembly of cytoskeletal proteins [44]. Supplementary figure 1 (Online Resource) shows the quantile-quantile (Q-Q) plots for each of the single cohort-specific EWAS.

Second, we meta-analysed the single cohort EWAS results, enabling an analysis of FTLD-associated differential cortical DNA methylation using tissue from 234 individuals. After a conservative Bonferroni adjustment for multiple testing (p<1.37 × 10^− 7^), the meta-analysis identified two differentially methylated CpGs in FTLD compared to controls, regardless of the pathological subgroup (FTLD-TDP or FTLD-tau), and corresponding subtypes (Fig. 3; Supplementary Fig. 2, Supplementary Table S2, Online Resource). The top CpG was annotated to a shore in the 5’UTR of *OTUD4* and was hypomethylated in FTLD compared to controls, whereas the other was annotated to a CpG island in the body of *NFATC1* and hypermethylated in FTLD compared to controls (Fig. 3). The direction of the effect was consistent across the three FTLD cohorts for these two hits, as well as for nine additional top meta-analysis loci obtained when considering a less stringent FDR p < 0.10 multiple testing correction (Fig. 3; Supplementary Table S2, Online Resource). Of note, none of these meta-analysis top differentially methylated sites showed epigenome-wide significant changes in FTLD3 alone (Supplementary Table S1, Online Resource) or in previous Alzheimer’s disease EWAS meta-analyses (Supplementary Table S2, Online Resource).

**Figure 3.**
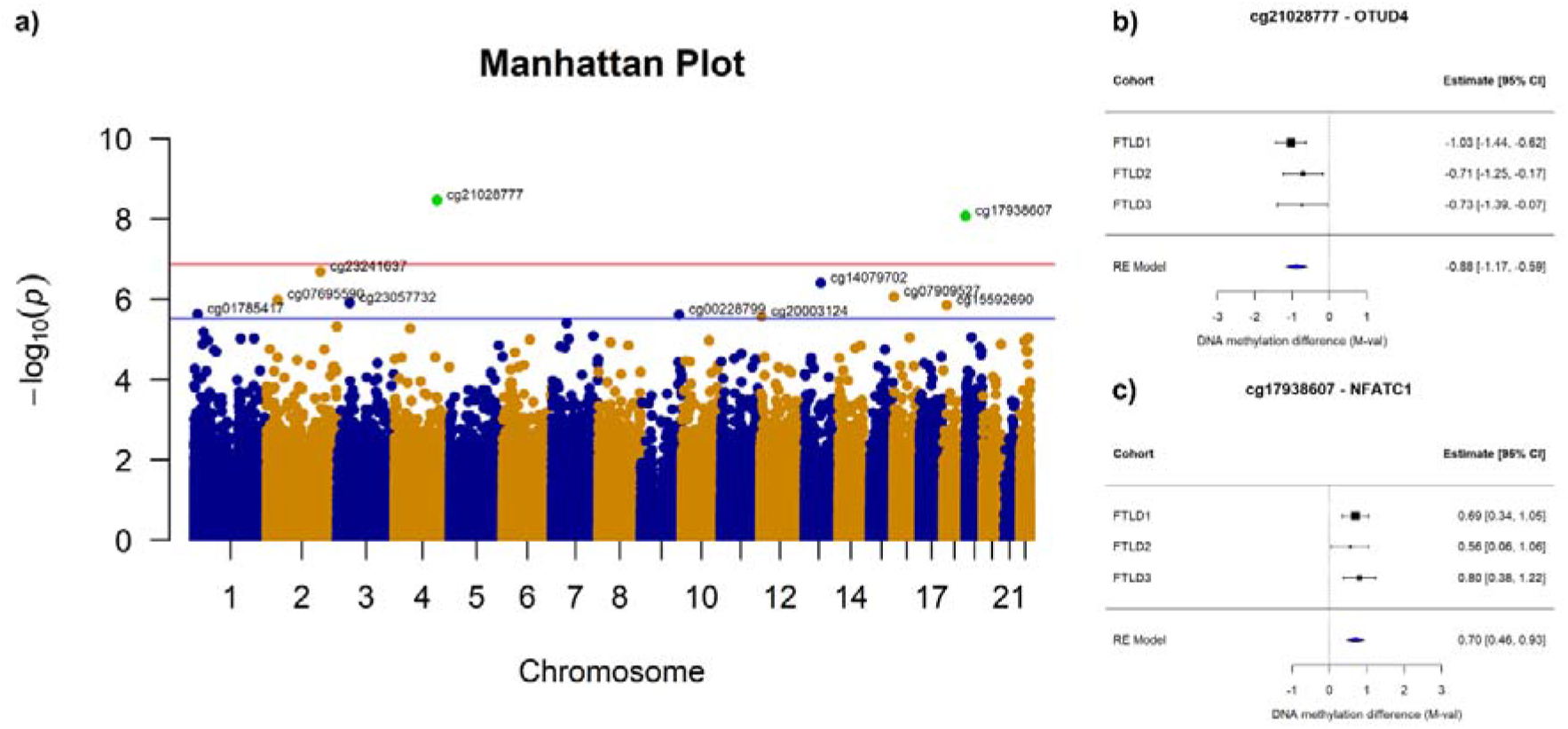
Differentially methylated positions identified in a case-control FTLD cross-cohort meta-analysis. **a)** Manhattan plot showing associations between single DNA methylation sites (CpGs) and FTLD from the meta-analysis random-effect results (total N = 234). CpGs are plotted on the x-axis according to their positions on each chromosome against association with FTLD on the y-axis (− log 10 p-value). The top red line indicates the conservative Bonferroni significance threshold (α) of p = 1.37 × 10^-7^. Green points indicate CpGs passing the Bonferroni threshold. The blue line indicates a less stringent threshold of p = 2.70 × 10^-6^ (FDR p = 0.10). **b)** Forest plot depicting the CpG in OTUD4, which is significantly hypomethylated in FTLD compared to controls in the cross-cohort meta-analysis (FTLD1 N=23, FTLD2 N=48, and FTLD3 N=163). c) Forest plot depicting the CpG in NFATC1, which is significantly hypermethylated in FTLD compared to controls in the cross-cohort meta-analysis (FTLD1 N=23, FTLD2 N=48, and FTLD3 N=163).

### Frontal cortex FTLD EWAS meta-analysis hits are consistent with downstream changes in mRNA and protein expression patterns

To investigate the downstream consequences of DNA methylation variation on gene expression in FTLD, we investigated available FTLD-TDP and FTLD-tau transcriptomic data [30, 82], as well as FTLD-TDP proteomics data. From the EWAS meta-analysis hits passing Bonferroni correction, consistent results were observed in both FTLD-TDP (frontal cortex) and FTLD-tau (temporal cortex) for OTUD4, which showed higher mRNA expression levels in FTLD cases compared to controls (Fig. 4). For *NFATC1*, increased expression was observed in FTLD-TDP when compared to controls (Fig. 4). However, this increase in expression was not observed in the FTLD-Tau, as seen in data from Wang et al. [82]. Of the nine additional top meta-analysis loci, *ZNF804A* showed lower mRNA expression levels and *IMPA2* showed higher mRNA expression levels in FTLD cases compared to controls (p < 0.05, Supplementary Fig. 3, Online Resource). DNA methylation levels in upstream regulatory regions are often inversely associated with gene expression levels [59, 79]. Therefore, lower methylation levels in CpGs annotated to 5’UTR in *OTUD4* and to TSS200 in *IMPA2*, and higher expression of these genes in FTLD compared to controls, meets such expectations. On the other hand, DNA methylation levels in gene bodies are usually positively associated with gene expression. Again, results align with this in the case of *NFATC1* (which showed higher methylation and higher expression in FTLD-TDP compared to controls) and *ZNF804A* (which showed lower methylation and lower expression in FTLD).

**Figure 4.**
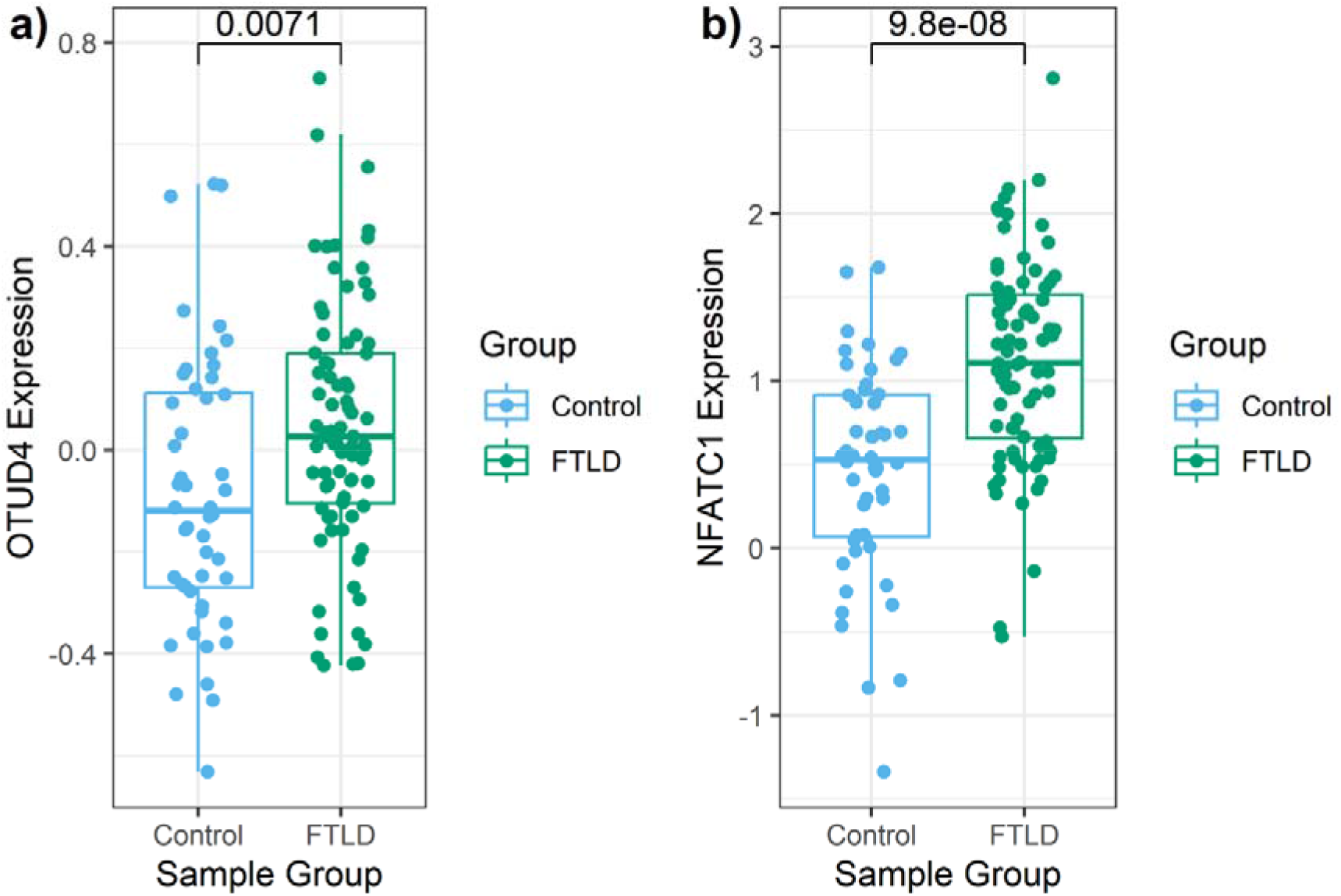
Boxplots showing gene expression levels for the two EWAS meta-analysis hits in FTLD-TDP and controls. RNA sequencing data from Hasan et al. [30] adjusted for age, sex, and RNA integrity number was used. Log2-transformed gene expression data is shown in the y-axis, and non-paired t-test p-value for the comparison between FTLD-TDP (N= 80) and controls (N= 48) is denoted at the top.

Only one of the two Bonferroni adjusted meta-analysis gene hits were detected in the frontal cortex proteomics data. *OTUD4* protein was upregulated in FTLD-TDP in types A and C compared to controls (Fig. 5), with the highest fold-change being observed in type C for the supernatant soluble fraction (fold-change = 14.72). These findings are in line with our observations with the RNAseq data and support consistent dysregulation of the *OTUD4* EWAS meta-analysis hit in FTLD. Therefore, we further investigated the patterns of OTUD4 protein expression in the frontal cortex and performed anti-OTUD4 immunohistochemical analysis (Fig. 6) using FTLD-TDP types A and C cases as well as controls that overlap with those used in the DNA methylation analysis (subset of the FTLD1 cohort). Minimal neuronal cytoplasmic staining was observed in the normal controls. However, in the FTLD-TDP cases, an increase in cytoplasmic staining intensity was observed in both the grey and white matter. In the grey matter, neuronal cytoplasmic staining was seen together with glial nuclear staining. In the white matter, there was an increase in glial staining. These results concur with the results from our proteomics and transcriptomics data.

**Figure 5.**
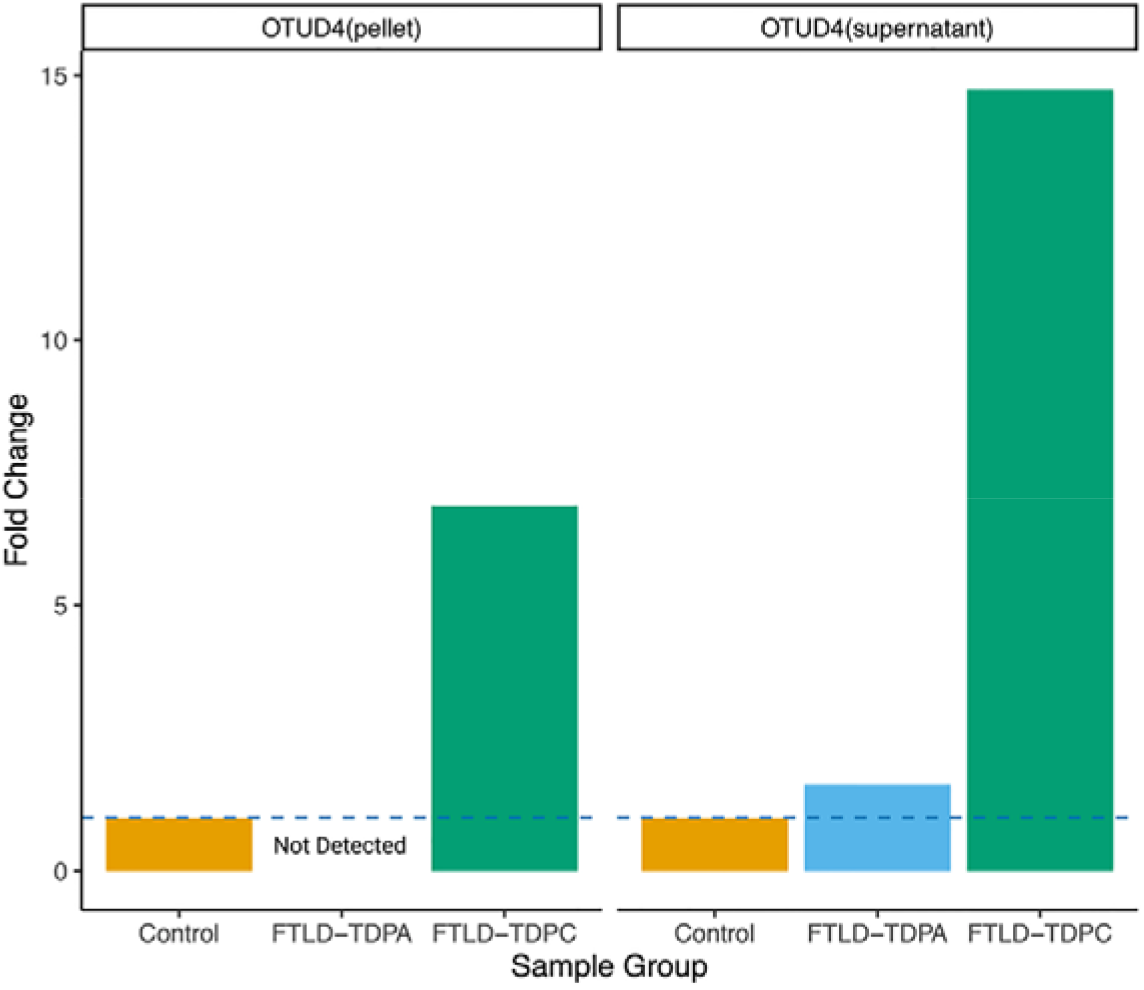
Bar plots of protein quantifications for the EWAS meta-analysis hit *OTUD4* in FTLD-TDP subtypes and controls. Out of the two EWAS meta-analysis hits, only the OTUD4 protein were detected in the proteomics data and are presented here. OTUD4 was detected in both fractions (pellet and supernatant). Two pooled samples (2 x 3 samples) per group were analysed. The average values were obtained for each group, and fold-changes were calculated comparing FTLD-TDP subtypes with controls. Bar plots show mean fold-change.

**Figure 6.**
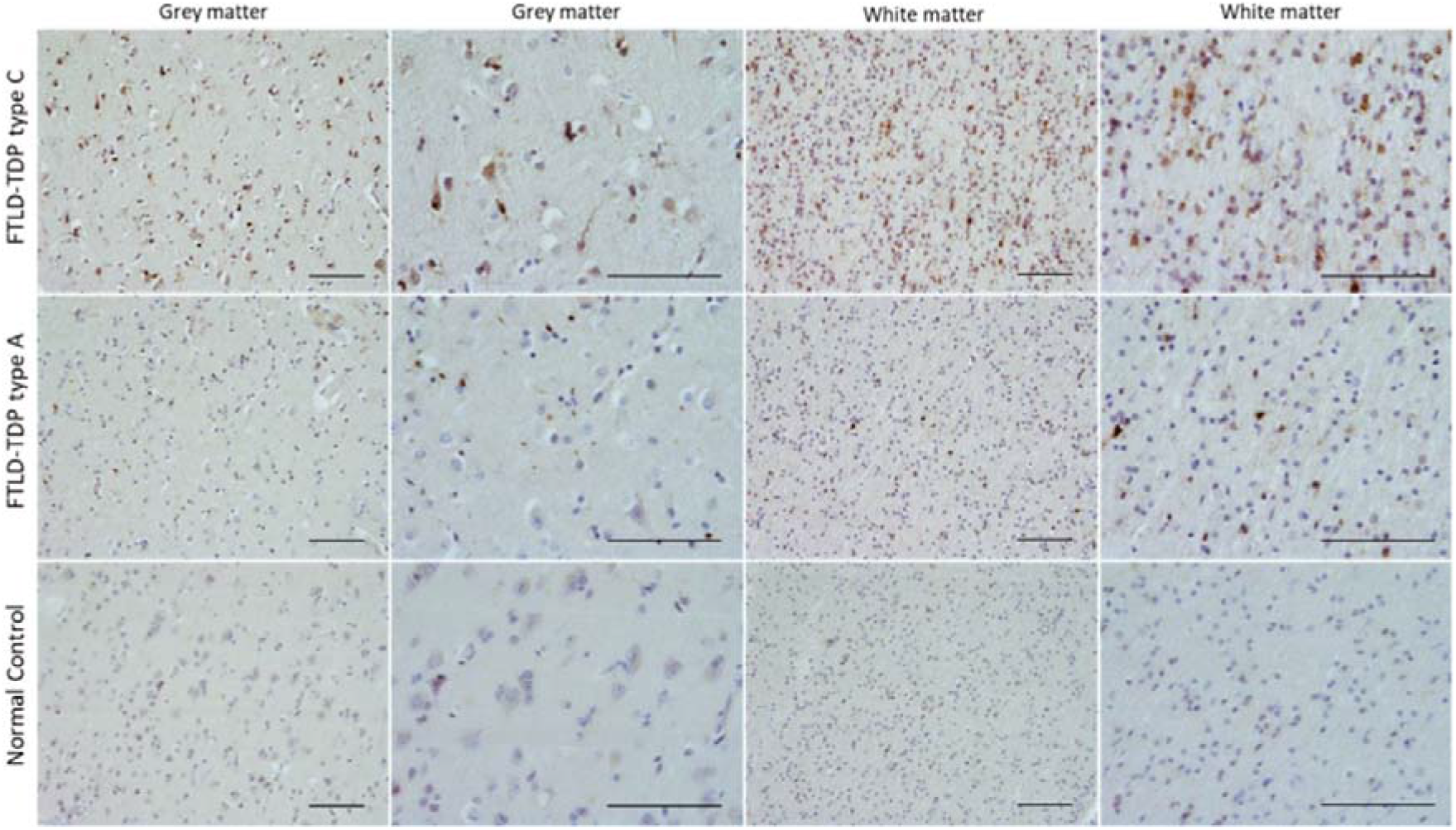
Immunoreactivity of *OTUD4* in FTLD-TDP (N=4 type A and N=3 type C) and controls (N=3). Immunohistochemical analysis was carried out in FFPE frontal cortex tissue from FTLD-TDP cases and controls overlapping with FTLD1, using a rabbit anti-OTUD4 antibody (Atlas Antibodies HPA036623, 1:200). Scale-bars represent 100 µm.

### DNA co-methylation modules are associated with the FTLD status, FTLD pathological subtypes, and disease-related traits

To provide insight into higher order relationships across DNA methylation sites (CpGs), we used an agnostic systems biology approach based on WGCNA and constructed co-methylation networks. Considering the top 20% most variable CpGs in each of the 3 cohorts (N = 56,001 CpGs), we identified clusters of highly correlated CpGs, henceforth called co-methylation modules, each assigned a colour name.

For the FTLD1, FTLD2 and FTLD3 networks, 9/33 (p<0.002, 0.05/33 modules), 16/49 (p<0.001, 0.05/49 modules) and 10/14 (p<0.004, 0.05/14 modules) co-methylation modules were found to be associated with the disease status (i.e., FTLD or control), respectively (Fig. 7a-c). Our co-methylation network analysis also revealed modules associated with specific pathological subgroup/subtypes in FTLD1 and FTLD2 networks (Supplementary Fig. 4, Online Resource). In a few cases, opposite effect directions were shown in one subgroup/subtype compared to another (e.g., midnightblue and salmon modules in FTLD1 TDPA vs TDPC, Supplementary Fig. 4a; and turquoise module in FTLD2 TDP vs Tau, Supplementary Fig. 4b; Online Resource). More detailed identification of subtype-specific DNA methylation signatures warrants further investigation in future studies.

**Figure 7.**
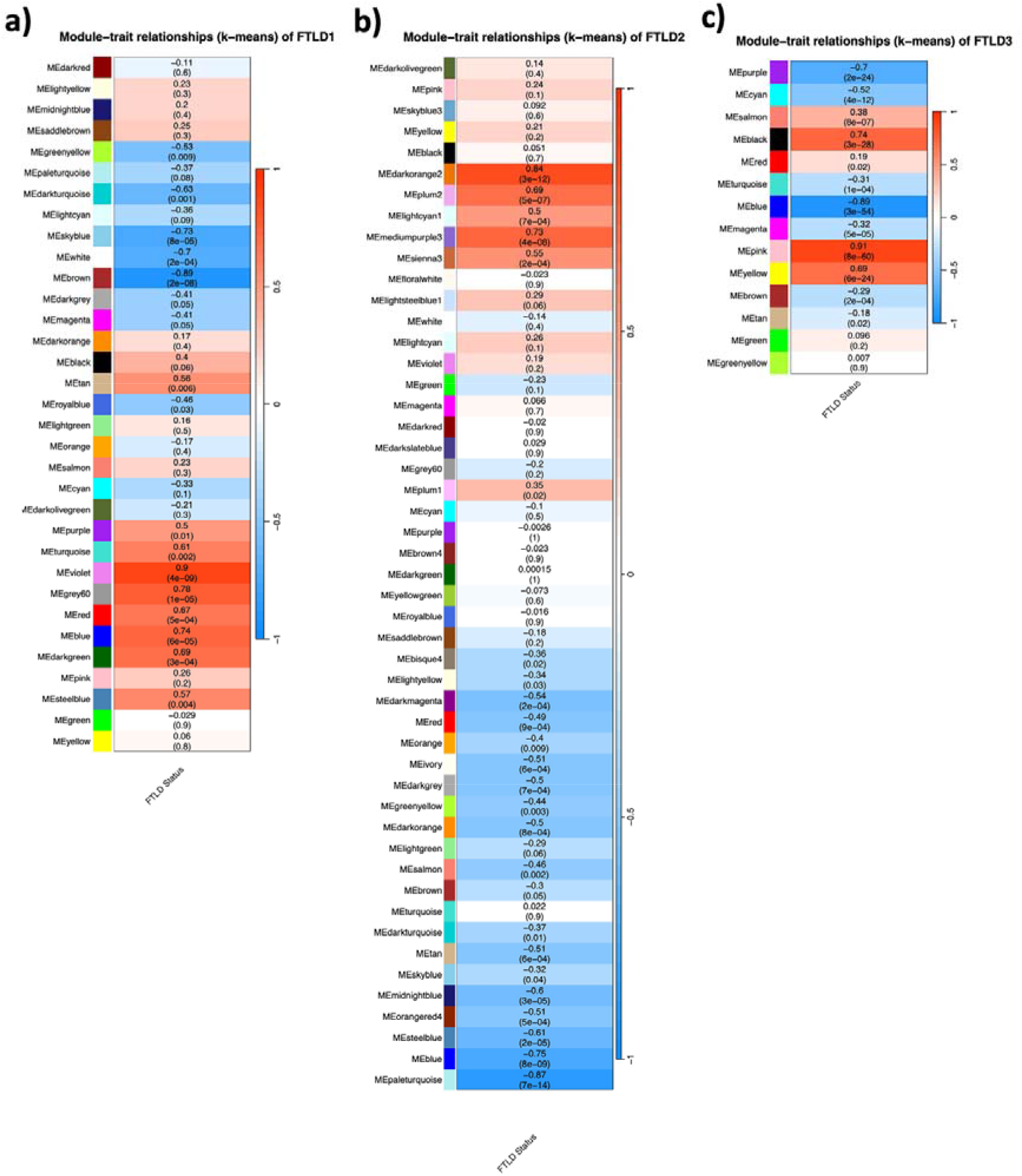
Module-trait correlations for the FTLD co-methylation networks. **a)** FTLD1; **b)** FTLD3; **c)** FTLD2. The rows represent the co-methylation module eigengenes (ME) and their colours, and the column represents the correlation of the methylation levels of CpGs in each module with the disease status. P-values are presented within each cell and the colour scale at the right indicates the strength of the correlation (darker cells depict stronger correlations, with blue representing negative and red representing positive correlations).

We also tested for correlations with additional disease-related traits as available for FTLD1, FTLD2, and FTLD3. We found associations between FTLD-associated co-methylation modules and disease duration as well as with macroscopic and/or microscopic measures of atrophy/neurodegeneration in the frontal and temporal lobes (Supplementary Fig. 4a-b, Online Resource). Two out of the ten modules associated with the disease status in FTLD3 were also associated with tau pathological burden (Braak stage, Supplementary Fig. 4c, Online Resource).

To assess replication of FTLD-associated co-methylation modules across datasets, we then ran preservation analysis for each dataset against each of the networks. We found that most of the FTLD-associated co-methylation modules were indeed moderately to highly preserved (Z-summary > 2) in at least one of the other two datasets (Supplementary Fig. 5, Online Resource), further supporting their relevance to FTLD regardless of the pathological subgroup/subtype. Exceptions to this were observed only for the FTLD1 brown, darkturquoise and grey60, and the FTLD2 darkorange2 modules, which seem to be perturbed in the other two datasets.

### Genes that compose FTLD-associated co-methylation modules are involved in transcription regulation, phosphorylation, the ubiquitin system and actin cytoskeleton dynamics

We then performed functional enrichment analysis to investigate which gene ontologies were shared across the three FTLD co-methylation networks. We found significant enrichment of terms related with transcription regulation (e.g., “DNA-binding transcription factor binding”), phosphorylation (“protein serine/threonine/tyrosine kinase activity”), the ubiquitin system (e.g., “ubiquitin protein ligase activity”), and actin cytoskeleton dynamics (e.g., “actin filament binding”). This was observed across the three co-methylation networks and across different modules of each network (Fig. 8). Dysregulation of all these processes had been previously linked to FTLD [68], and our findings now support a role for DNA methylation as a mechanism involved in such dysregulation.

**Figure 8.**
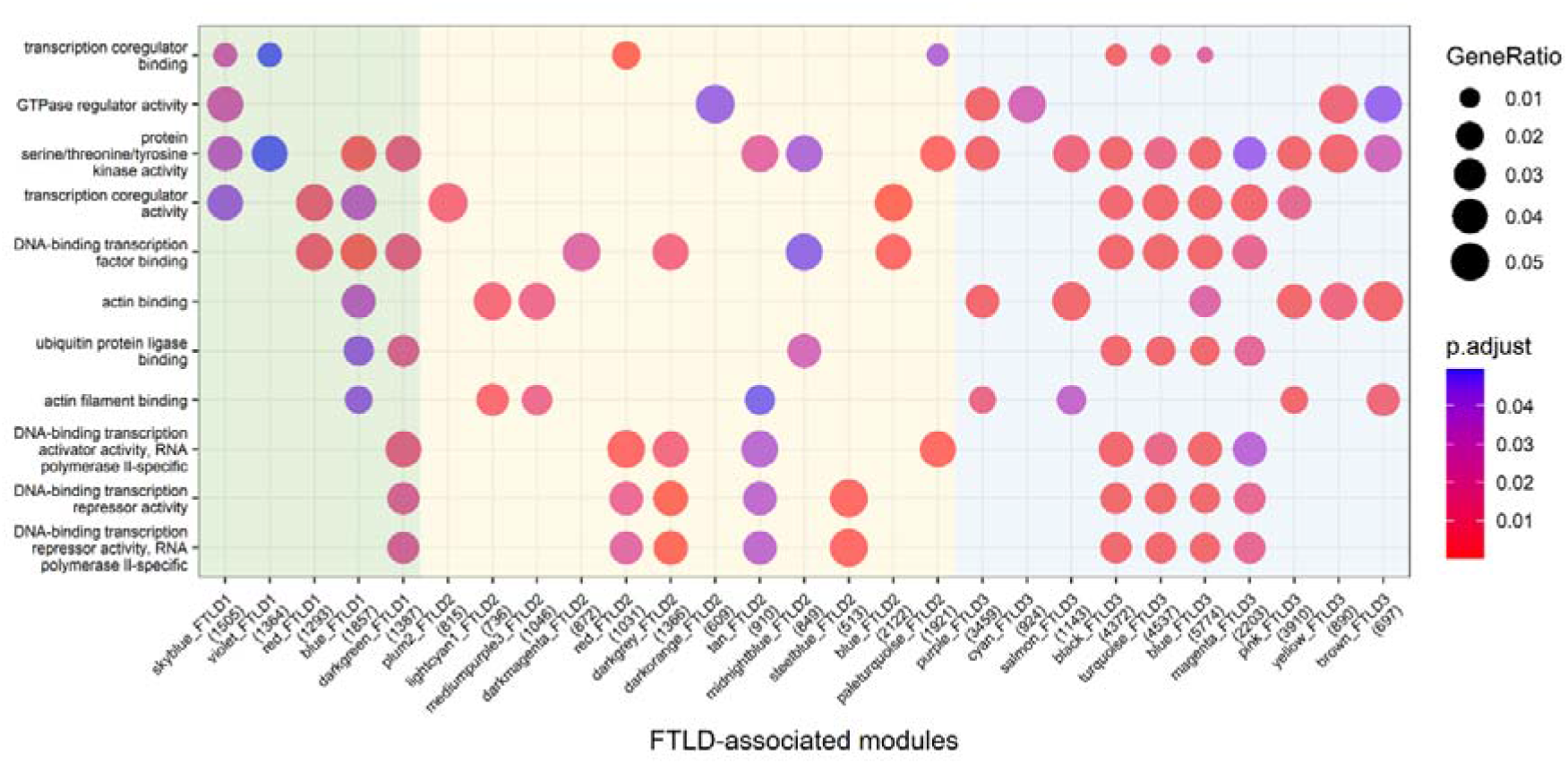
Functional enrichment for the FTLD-associated co-methylation modules across the three networks. Y-axis shows top enriched gene ontology terms, while x-axis depicts FTLD-associated modules in FTLD1 (green), FTLD2 (yellow) and FTLD3 (blue) co-methylation networks. Modules not showing enrichment for shared terms across the networks are not shown.

### FTLD-associated modules are enriched for genes relevant for pyramidal neurons and endothelial cells across all three co-methylation networks

We also aimed to elucidate whether the genes that compose FTLD-associated co-methylation modules are relevant for specific brain cell-types. Across the three networks (FTLD1, FTLD2 and FTLD3), we found significant enrichments for pyramidal neurons and endothelial/mural cells (Fig. 9), suggesting these cell types are consistently affected by the DNA methylation changes in FTLD regardless of the pathological subgroup/subtype. Previous studies with pathological assessment, as well as transcriptomic analysis in FTLD brain tissue, support changes in these cell types in FTLD [24–26, 30, 63]. Additionally, in the FTLD1 and the FTLD3 networks, we found signatures with an overrepresentation of oligodendrocyte markers. Of note, FTLD3 is composed of PSP cases, which, unlike the other FTLD groups studied here, is known to present with pathological accumulation of tau in the oligodendrocytes [84]. The FTLD3 network was also enriched for microglia and interneurons.

**Figure 9.**
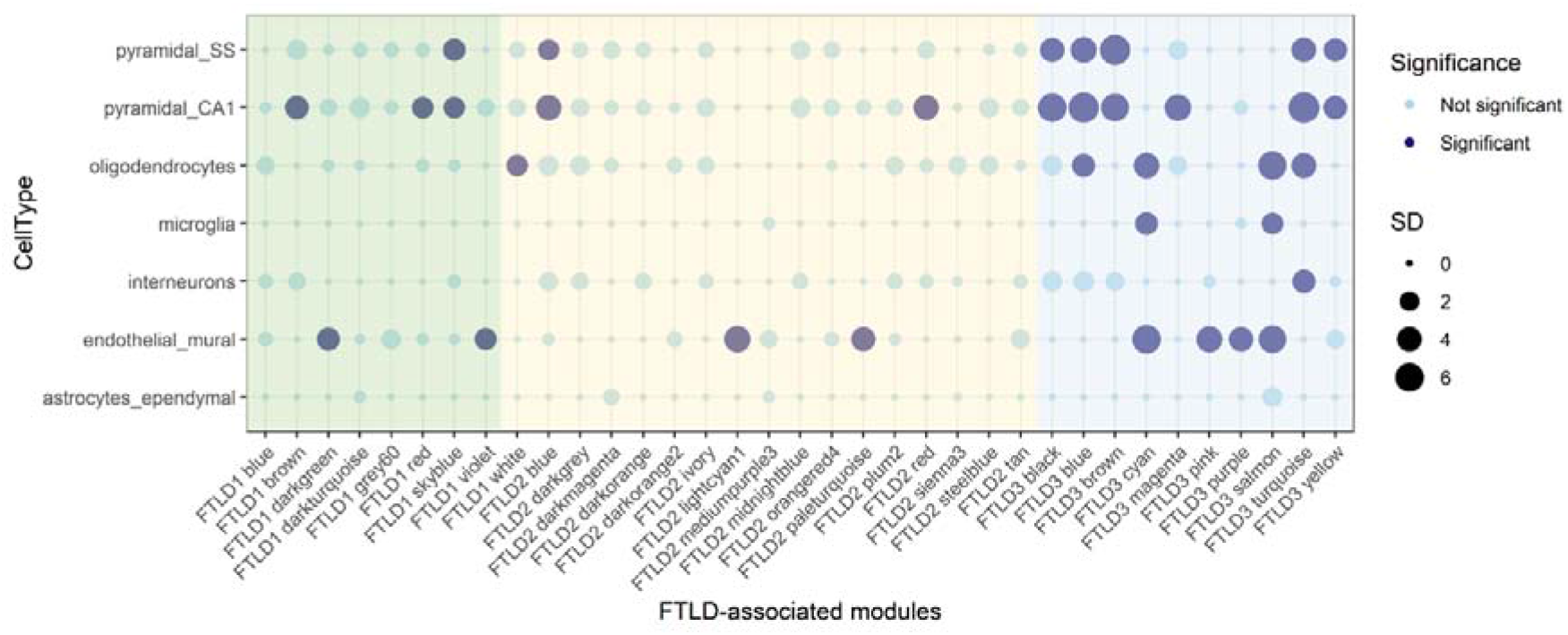
Cell-type enrichment for all FTLD-associated co-methylation modules across the three co-methylation networks. Green denotes FTLD-associated modules in the FTLD1 network; Yellow denotes FTLD-associated modules in the FTLD2 network; Blue denotes FTLD-associated modules in the FTLD3 network. Dark filled circles highlight the cell types found to be significantly enriched with adjusted p < 0.05 after Bonferroni correction over all cell types within each module; the size of the circles represents the number of standard deviations (SD) from the mean. Cell-type enrichment analysis on the FTLD-related modules was performed using the package EWCE [73] and associated single-cell transcriptomic data [88].

### OTUD4 and other top meta-analysis loci are co-methylated in all three networks

We then examined whether the 11 EWAS meta-analysis top loci (FDR p < 0.10) were present (Supplementary Table S2, Online Resource) in the co-methylation networks and whether any co-methylation modules were enriched for such loci (Supplementary Table S3, Online Resource). Notably, the top meta-analysis hit in *OTUD4* was present in all three networks (FTLD1 – brown, FTLD2 – blue, and FTLD3 – blue modules), and was always co-methylated with the CpG annotated to CEBPZ (Supplementary Table S3, Online Resource). These modules showed a significant enrichment for the top EWAS meta-analysis loci [Fisher’s exact test, FTLD1 – brown odds ratio (OR) = 14.9, p = 0.003; FTLD2 – blue OR = 10.6, p = 0.007; FTLD3 – blue OR = 8.0, p = 0.017). We therefore decided to further investigate similarities across these three modules (FTLD1 – brown, FTLD2 – blue, and FTLD3 – blue), which will henceforth be referred to as “OTUD4-modules”.

It is of note that only eight CpGs were shared across the three “*OTUD4*-modules”, two of which – cg21028777 in *OTUD4* and cg07695590 in *CEBPZ* – correspond to top EWAS meta-analysis loci (Supplementary Fig. 6, Online Resource), highlighting their importance across the FTLD subgroups/subtypes. All three “OTUD4-modules” were inversely related with the disease status, i.e., lower levels of methylation in CpGs composing these modules are associated with increased risk of FTLD (Fig. 7; FTLD1 – brown r=-0.89, p = 2×10^-8^; FTLD2 – blue r=-0.75, p = 8×10^-9^; and FTLD3 – blue r=-0.89, p = 3×10^-54^). FTLD2 blue was also inversely associated with the severity of neuronal loss in the frontal cortex (r=-0.48, p=0.001, Supplementary Fig. 4b, Online Resource). Although not reaching statistical significance after accounting for multiple testing corrections, a similar trend was observed with the severity of neuronal loss in the temporal cortex for FTLD2 blue (r=-0.46, Supplementary Fig. 4b, Online Resource) as well as for FTLD1 brown in both frontal and temporal cortices (r=-0.29 n.s., and r=-0.63 p = 0.001, respectively, Supplementary Fig. 4a, Online Resource). These findings further support the relevance of these signatures enriched for top EWAS meta-analysis loci, including CpGs in *OTUD4* and *CEBPZ*, in disease progression/severity.

Previous studies have shown that *OTUD4* [20], tau [5], TDP-43 and a growing number of additional FTLD-related RNA-binding proteins [10] play an important role in the biology of stress granules. We therefore investigated whether stress granules proteins and *OTUD4* protein interactors were present in the “OTUD4-modules”. Indeed, many genes encoding for such proteins were represented in these modules, including several genes associated with genetic FTLD risk such as *MAPT* (encoding for tau), present across the three “OTUD4-modules”, and *FUS*, present in FTLD3-blue (Supplementary Tables S4 and S5, Online Resource). The same was true for many hnRNPs, such as *HNRNPA1, HNRNPC*, and *HNRNPUL1*, which are present in the “OTUD4-modules” and are OTUD4 protein interactors (Supplementary Tables S4 and S5, Online Resource). These hnRNPs are also known targets of the transcription factor *CEBPZ* (as described by Ma’ayan et al. [65]), which is also a top EWAS meta-analysis loci and is co-methylated with *OTUD4* across the networks.

We also identified the hub genes in the three “OTUD4-modules” (i.e., the most interconnected genes within the module). These were *ADCY1, TLE6* and *GDAP1* for FTLD1-brown, FTLD2-blue and FTLD3-blue, respectively (Supplementary Table S4, Online Resource). Of note and highly relevant for FTLD, *ADCY1* has been found to be implicated in learning, memory, and behaviour [71]. The importance of TLE6 to brain related disease is supported through its association with bipolar disorder [21], and mutations in *GDAP1* cause inherited peripheral neuropathies [61].

### “*OTUD4*-modules” implicate glutamatergic synapse and pyramidal neurons

More detailed gene ontology enrichment of “OTUD4-modules” once again highlighted transcriptional regulation and the ubiquitin system, as well as nuclear speck, synapse (particularly glutamatergic synapse), and axon development (Supplementary Fig. 7, Online Resource). All three meta-hit modules showed an enrichment for pyramidal neurons and the FTLD3 blue module additionally showed an enrichment for oligodendrocytes (Fig. 9). Further supporting the importance of *OTUD4* and *CEBPZ* in glutamatergic cells, in the normal brain (human and mouse) these genes show the highest expression in glutamatergic neurons and/or cortical and hippocampal pyramidal and granule cell layers (Supplementary Fig. 8-9, Online Resource).

Using gene expression data and derived cellular proportions from Hasan et al. [30], we observed a positive relationship between both *OTUD4* and *CEBPZ* expression and proportions of excitatory neurons in controls and FTLD-TDP type A (Supplementary Fig. 10, Online Resource). This finding further supports the relevance of *OTUD4* and *CEBPZ* in excitatory glutamatergic neurons. However, that relationship is perturbed in FTLD-TDP type C (Supplementary Fig. 10, Online Resource), which could suggest higher expression of these genes by fewer surviving excitatory neurons and/or higher expression by other cell type(s).

## DISCUSSION

We have conducted,to our knowledge, the first FTLD EWAS meta-analysis utilizing three independent cohorts and incorporating results from 234 brain donors. We identified two differentially methylated CpGs shared across a range of FTLD subgroups (FTLD-TDP and FTLD-tau) and corresponding subtypes, which map to *OTUD4* and *NFATC1*. Systems biology approaches such as co-methylation network analysis are powerful methodologies for identifying pathways and networks which may be more relevant to disease pathophysiology than individual genes. We therefore performed a co-methylation network analysis in each of the independent cohorts and identified modules associated with the FTLD disease status and FTLD-related traits. Interestingly, *CEBPZ* always clustered with OTUD4, and the “OTUD4-modules” were enriched for meta-analysis top loci in each of the three independent cohorts. Using functional and cell-type enrichment analysis of modules of interest, we identified several biological processes with relevance to FTLD pathology, including the ubiquitin system, RNA granule formation and glutamatergic synaptic signalling, which we discuss below. It is of note that none of the loci identified in our meta-analysis match with neuropathology-associated loci identified in large AD studies [72, 74, 89], therefore supporting the hypothesis that molecular changes in these loci reflect shared disease biology aspects of FTLD subgroups/subtypes rather than a mere downstream consequence of neurodegeneration.

The *OTUD4* gene encodes the protein OTUD domain-containing protein 4, a de-ubiquitinating enzyme [54]. Mutations in this gene are associated with Gordon Holmes syndrome, which is characterised by ataxia and hypogonadotropism [48]. Interestingly, a combination of mutations in *OTUD4* along with mutations in *RNF216*, which codes for a ubiquitin ligase, was also found to result in dementia [48]. The protein is known to have roles in modulating inflammatory signalling [91] and in the alkylation damage response [90], and has more recently been demonstrated to interact with RNA binding proteins (RBPs), including TDP-43 (which aggregates in FTLD-TDP), which are important in the functioning of neuronal RNA granules and stress granules [18]. RNA granules are structures which facilitate the translocation and storage of mRNAs [37], whilst stress granules are formed when cellular stressors such as oxidative stress are present, possibly as a mechanism to reversibly block translation initiation until the stress has been removed [16, 35]. Notably, similarly to TDP-43 [6], *OTUD4* was shown to be important in the correct formation of stress granules [20]. Indeed, there is much evidence as to the importance of the ubiquitin system in the functioning of stress granules [36, 55, 77]. The hypomethylation of the 5’UTR region of *OTUD4* (cg21028777), which was observed as the top hit from the FTLD EWAS meta-analysis, and the inclusion of this CpG in three modules where decreased methylation was associated with increased risk of FTLD indicates that decreased methylation of this gene might be involved in the pathogenesis of FTLD. Further supporting these findings, the *OTUD4* gene and protein expression levels are dysregulated in FTLD [30, 82].

Also supporting the importance of the role of ubiquitination and granule formation are the results from the functional enrichment analysis of the three network modules containing OTUD4, which revealed an overrepresentation of terms relating to the ubiquitin system. All three meta-hit modules contained terms such as “ubiquitin protein ligase activity”, the FTLD2-blue module also showed enrichment of the GO term “ribonucleoprotein granule”, indicating that other genes in this module might also have processes relevant to granule formation, as with the meta-hit OTUD4. Ubiquitin signalling is well described as a process implicated in neurodegenerative disease pathology, and several genes involved in ubiquitin and ubiquitin binding processes are known to be mutated/contain risk alleles in multiple neurodegenerative diseases, including FTD [68].

Ontology terms enriched in our functional analysis of FTLD-associated modules also include many relating to regulation of transcription such as “DNA-binding transcription factor binding” and “transcription coregulator activity”. Another meta-analysis top loci was annotated to the *CEBPZ* gene, which encodes the CCAAT Enhancer Binding Protein Zeta, a transcription factor implicated in cellular response to environmental stimuli through transcriptional processes that regulate heat shock factors, including HSP70 [47]. HSP70 is a heat-shock protein involved in several protein folding processes, including the refolding of aggregated proteins [32, 46, 62]. Furthermore, HSP70 has been shown to have a role in the prevention of build-up of misfolded proteins in stress granules [49]. Interestingly, a CpG in *PFDN6* was the top-most differentially methylated CpG in the FTLD3 (FTLD-tau) EWAS. This gene encodes for the subunit 6 of prefoldin, which is a co-chaperone of HSP70, regulates the correct folding of proteins and is involved in the proper assembly of cytoskeletal proteins [44]. Prefoldin proteins themselves have also been associated with neurodegenerative disease pathology [44, 75].

Our functional enrichment analysis of the “OTUD4-modules”, FTLD1-brown, FTLD2-blue and FTLD3-blue, showed that these modules were enriched for GO terms (for cellular component) relating to synapses, including “synaptic membrane”, “asymmetric synapse”, “postsynaptic density”, and “glutamatergic synapse”. Cell-type enrichment analysis revealed that these three modules were also significantly enriched for markers of pyramidal/glutamatergic cells. These findings were further substantiated with expression patterns of *OTUD4* and *CEBPZ* single-nuclei and mouse expression data. Glutamate, which is the most abundant excitatory neurotransmitter in the human brain [92], is typically associated with memory, learning and other higher cognitive functions [12], and has also been implicated in neurodegeneration [58]. The contribution of neurotransmitter deficits, and specifically, changes in glutamate and glutamate signalling have been described in FTD [2, 14, 29, 33, 56]. DNA methylation has previously been suggested to be an important regulator of glutamatergic synaptic scaling (also known as homeostatic synaptic plasticity), with demethylation found to be associated with increased glutamatergic synapse strength in cultured neurons [52], we here find evidence supporting disruption of such processes in FTLD. Homeostatic synaptic plasticity has been linked to neurodegeneration, possibly with loss of function due to pathogenesis, or through an increase as a mechanism to preserve function despite neurodegenerative deficits [60]. There is a known link between RNA granule formation and synapse plasticity; with RNA-binding protein function known to be particularly important. This has been proposed to be dysregulated in FTLD, whereby mutations in the genes encoding for TDP-43 and FUS lead to dysregulated granule formation dynamics and consequent disturbances in mRNA translation and synaptic function [45, 70]. Moreover, the levels of known *OTUD4* protein interactor FMRP are regulated by ubiquitination in response to stimulation by the metabotropic glutamate receptor [31, 57], and this is involved in the regulation of synaptic plasticity, providing another possible link between separate findings in our study.

The *NFATC1* gene, which was also identified as an FTLD-associated loci in the EWAS meta-analysis, encodes the nuclear factor of activated T cells 1, and belongs to the NFAT family of activity-dependent transcription factors. In the nervous system, the NFAT family has been shown to play a regulatory role in neuronal excitability, axonal growth, synaptic plasticity, and neuronal survival [80]. Aberrant NFAT-related signalling has been reported in AD, and NFAT1 seems to be selectively activated early in cognitive decline [1], supporting its possible involvement in the pathogenesis of neurodegenerative diseases/dementias.

As is the case with any other genome-wide DNA methylation study, there are key limitations. First, by studying post-mortem tissue, i.e., the end stage of the disease, causality cannot be elucidated. Second, because FTLD is heterogeneous, comprising several pathological subgroups and subtypes, and given the relatively small sample size per subtype, this might have hampered the identification of additional DNA methylation alterations, especially subtype-specific variation. Notwithstanding, we focused on the shared DNA methylation variation across FTLD subgroups/subtypes, and we used independent and complementary analytical approaches (EWAS followed by meta-analysis, and co-methylation network analysis followed by preservation analysis) and datasets, which identified concordant results and consistently identified the involvement of *OTUD4* and related genes in FTLD.

In summary, this study identified genome-wide DNA methylation changes in post-mortem frontal cortex tissue of FTLD subjects, highlighting new FTLD-associated loci, and implicated DNA methylation as a mechanism involved in the dysregulation of important processes such as ubiquitin and glutamatergic signalling in FTLD. Our findings increase the understanding of the biology of FTLD and role of DNA methylation its pathophysiology, pointing towards new avenues that could be explored for therapeutic development.

## Acknowledgements

The authors would like to thank UCL Genomics centre for advice and processing of the EPIC arrays for the FTLD1 cohort. The authors would also like to acknowledge the Queen Square Brain Bank (London, UK), and the Dutch Brain Bank, Netherlands Institute for Neuroscience (Amsterdam, Netherlands) for providing brain tissues from FTLD cases and controls. The Queen Square Brain Bank is supported by the Reta Lila Weston Institute of Neurological Studies, UCL Queen Square Institute of Neurology. KF is supported by the Medical Research Council (MR/N013867/1). MM is supported by a grant from the Multiple System Atrophy Trust awarded to CB. CB is supported by Alzheimer’s Research UK (ARUK-RF2019B-005) and the Multiple System Atrophy Trust. JH, RH, and TR are supported by NIH NIA R56-AG055824 and U01-AG068880, and NIH NINDS U54NS123743.

