## supplementary figures for "Brain DNA methylomic analysis of frontotemporal lobar degeneration reveals *OTUD4* in shared dysregulated signatures across pathological subtypes"

Department of Neurodegenerative Disease

UCL Queen Square Institute of Neurology

United Kingdom

**a)**

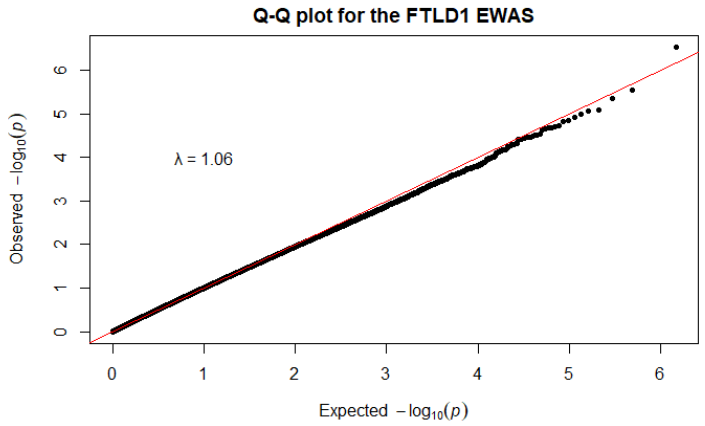


**b)**

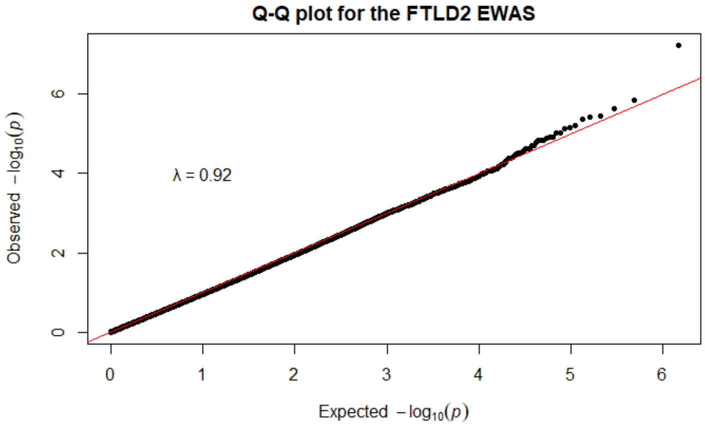


**c)**

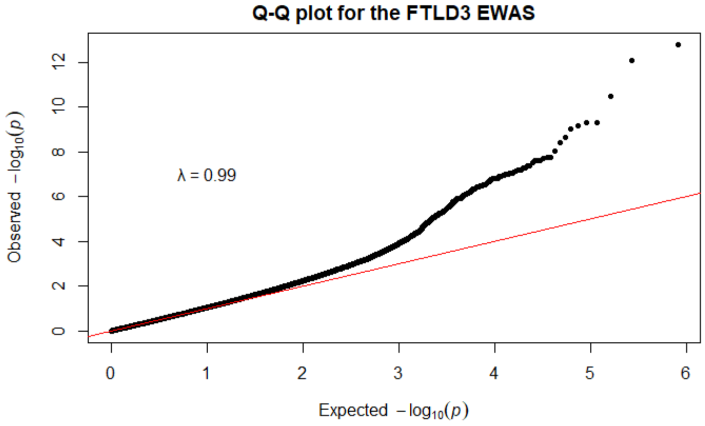


**Supplementary figure 1. Quantile-quantile (Q-Q) plots for the FTLD cohort-specific case-control EWAS. a)** Q-Q plot for the FTLD1 EWAS, with an estimated inflation factor (λ) of 1.06; **b)** Q-Q plot for the FTLD2 EWAS, with an estimated inflation factor (λ) of 0.92; **c)** Q-Q plot for the FTLD3 EWAS, with an estimated inflation factor (λ) of 0.99.

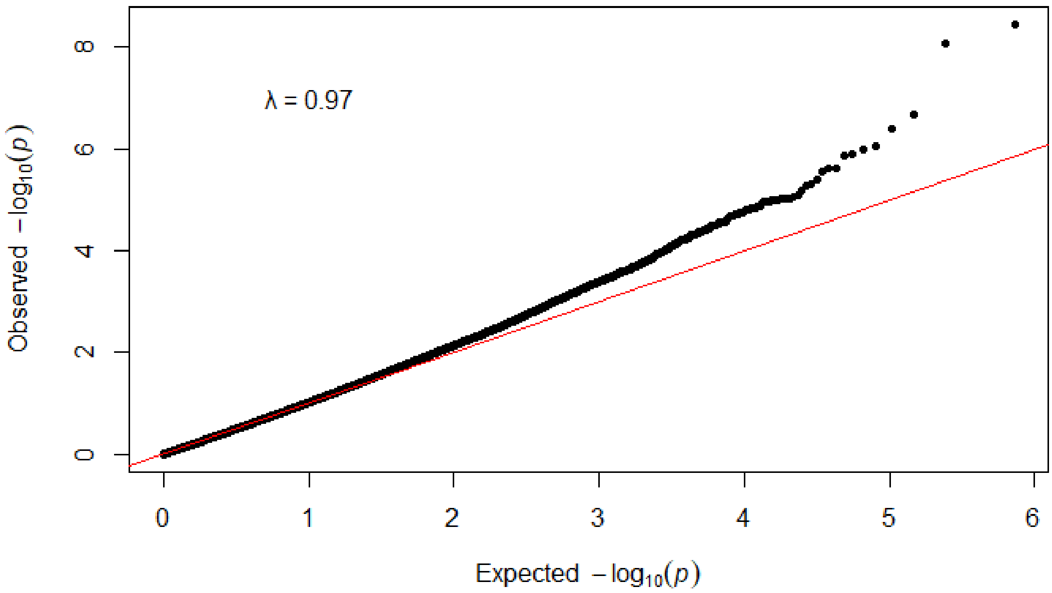


**Supplementary figure 2. Quantile-quantile (Q-Q) plots for the FTLD case-control EWAS meta-analysis with random-effect models across three cohorts (FTLD1, FTLD2 and FTLD3).**

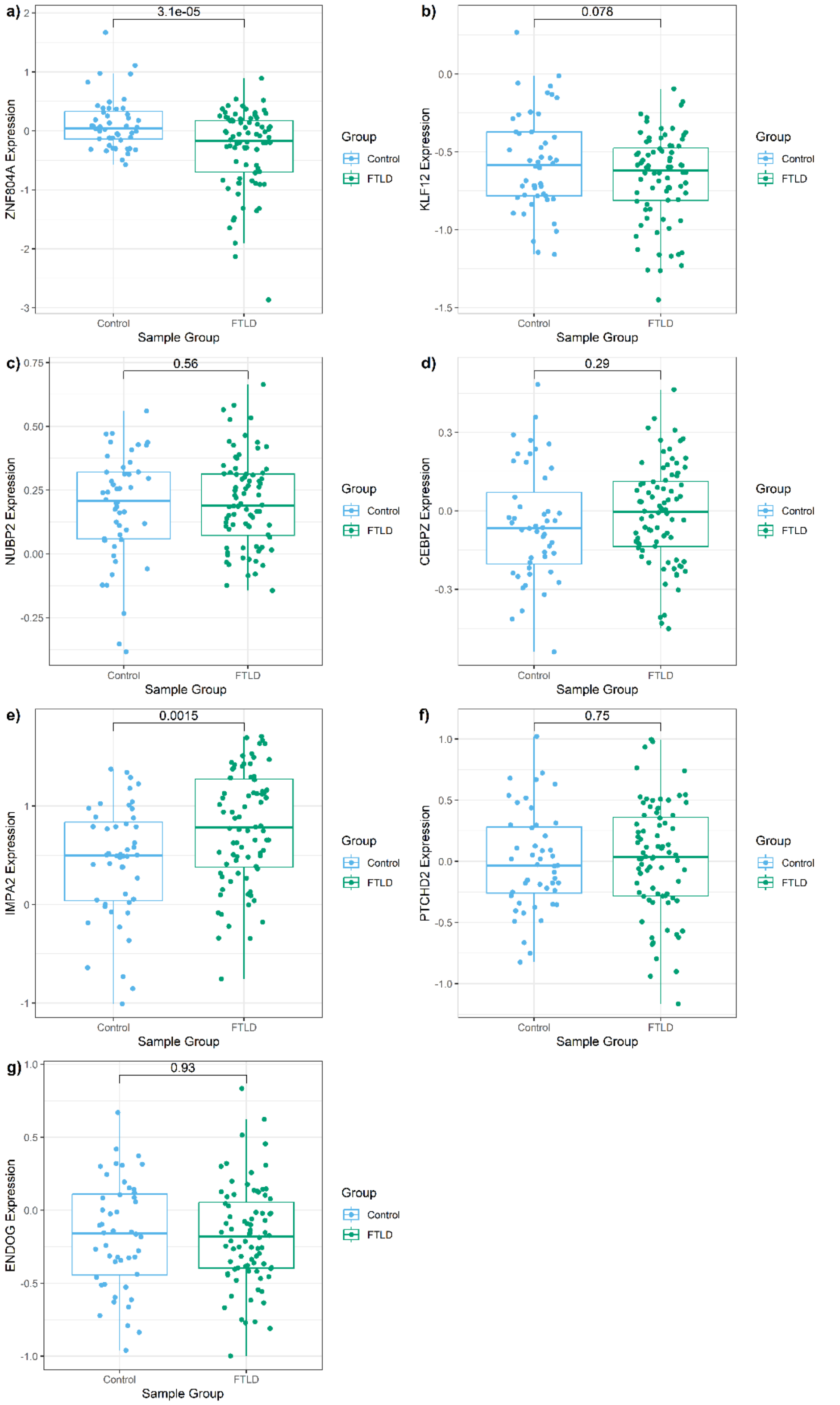


**Supplementary figure 3. Boxplots showing gene expression levels for the top EWAS meta-analysis loci in FTLD-TDP and controls.** RNA sequencing data from Hasan et al.[2] adjusted for age, sex, and RNA integrity number was used. Log2-transformed gene expression data is shown in the y-axis, and non-paired t-test p-value for the comparison between FTLD-TDP (N=80) and controls (N=48) is denoted at the top.

a) **Module−trait relationships (k−means) of FTLD1**

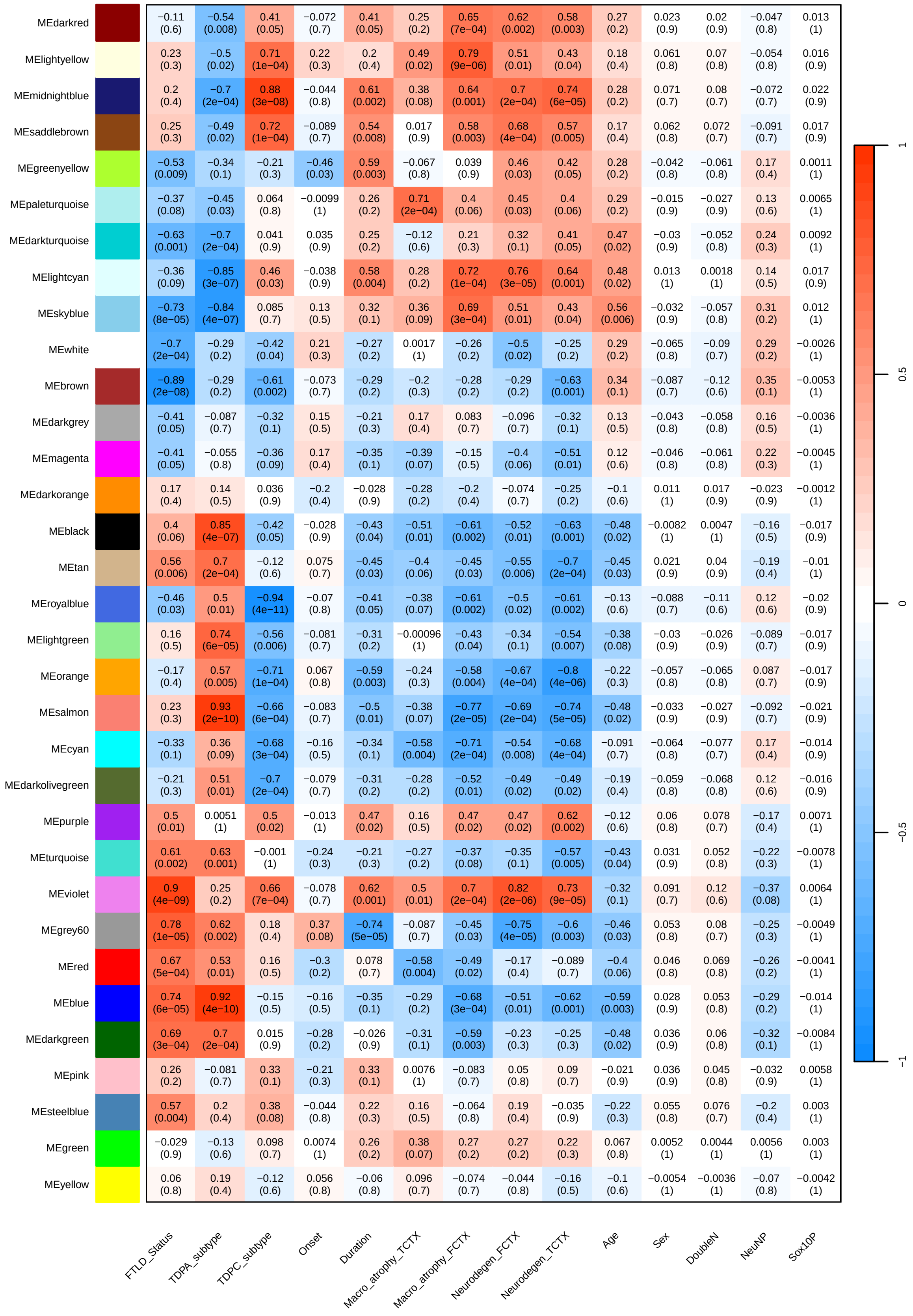

b)

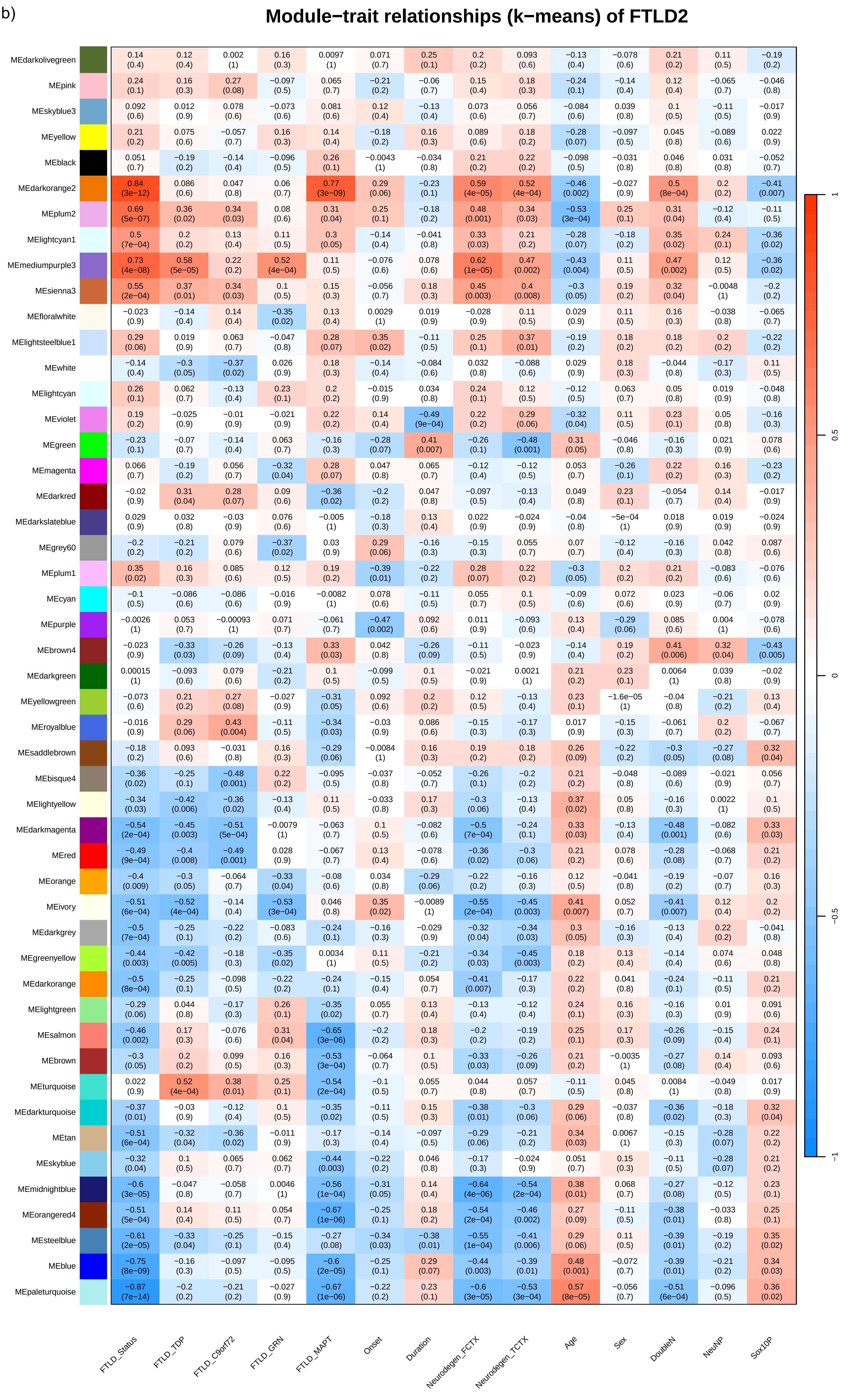

Module−trait relationships (k−means) of FTLD2
MEdarkolivegreen
MEpink
MEskyblue3
MEyellow
MEblack
MEdarkorange2
MEplum2
MElightcyan1
MEmediumpurple3
MEsienna3
MEfloralwhite
MElightsteelblue1
MEwhite
MElightcyan
MEviolet
MEgreen
MEmagenta
MEdarkred
MEdarkslateblue
MEgrey60
MEplum1
MEcyan
MEpurple
MEbrown4
MEdarkgreen
MEyellowgreen
MEroyalblue
MEsaddlebrown
MEbisque4
MElightyellow
MEdarkmagenta
MEred
MEorange
MEivory
MEdarkgrey
MEgreenyellow
MEdarkorange
MElightgreen
MEsalmon
MEbrown
MEturquoise
MEdarkturquoise
MEtan
MEskyblue
MEmidnightblue
MEorangered4
MEsteelblue
MEblue
MEpaleturquoise
FTLD_Status
FTLD_TDP
FTLD_C9orf72
FTLD_GRN
FTLD_MAPT
Onset
Duration
Neurodegen_FCTX
Neurodegen_TCTX
Age
Sex
DoubleN
NeuNP
Sox10P
1
0.5
0
−0.5
−1

c) **Module−trait relationships (k−means) of FTLD3**

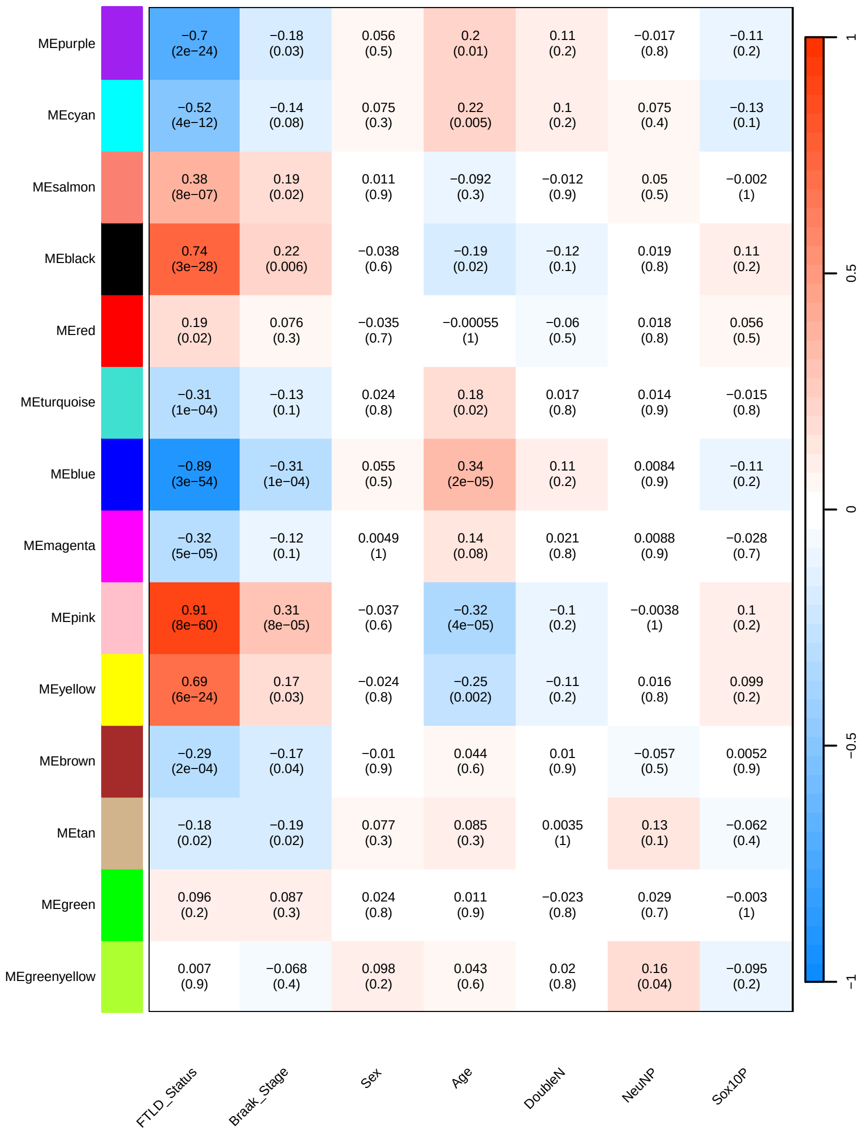

**Supplementary figure 4. Module-trait correlations for the FTLD co-methylation networks.** a) FTLD1; b) FTLD3; c) FTLD2. The rows represent the co-methylation module eigengene (ME) and its colour, and the columns represent clinical/pathological traits. P values are presented within each cell and the colour scale at the right indicates the strength of the correlation (darker cells depict stronger correlations, with blue representing negative and red positive correlations).

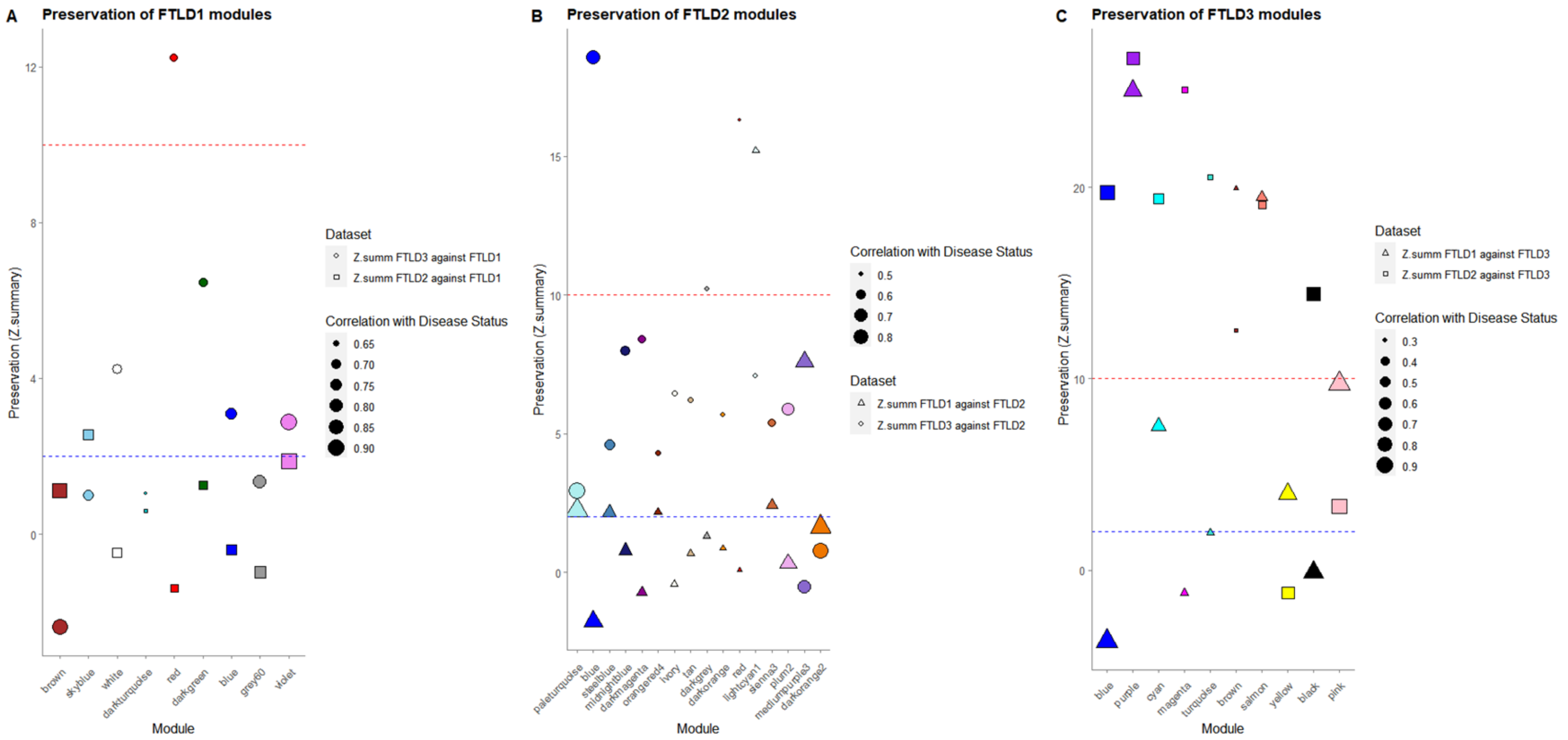


**Supplementary figure 5. Preservation analysis Z-summary results for the co-methylation modules associated with FTLD in each individual network. a)** Preservation of the FTLD1 network FTLD-associated modules in the FTLD2 (squares) and FTLD3 (circles) data; **b)** Preservation of the FTLD2 network FTLD-associated modules in the FTLD1 (triangles) and FTLD3 (circles) data; **c)** Preservation of the FTLD3 network FTLD-associated modules in the FTLD1 (triangles) and FTLD2 (squares) data. Colour symbols represent co-methylation modules associated with the FTLD status in the corresponding network, and the size of the symbols represents the magnitude of the association. Modules on the left hand side of the plot show the strongest negative associations with the FTLD status, while those on the right show the strongest positive associations with the FTLD status. Y-axis represents the preservation Z.summary. Modules above the red dashed line (Z.summary >10) are said to be highly preserved, modules between the blue and red dashed lines (2< Z.summary <10) are said to be moderately preserved. Modules under the blue dashed line (Z.summary <2) are not preserved.

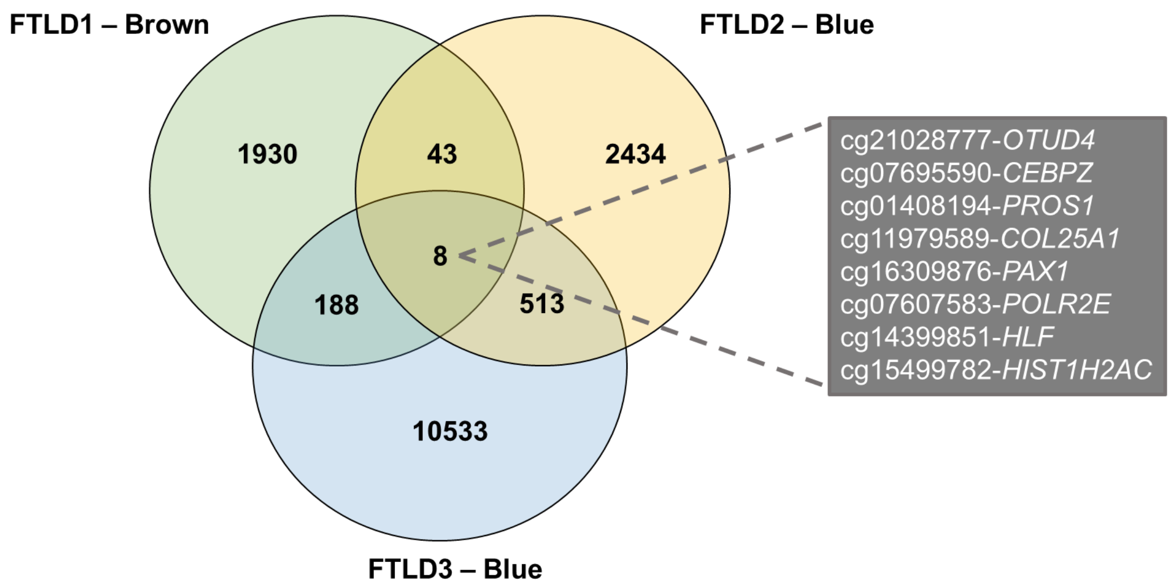


**Supplementary figure 6. CpG overlap across the FTLD meta-hit enriched modules.** From the three independent co-methylation networks three modules (FTLD1 – Brown, FTLD2 – Blue and FTLD3 – Blue) were enriched for the FTLD top EWAS meta-analysis hits. The eight probes that are present in all three modules are highlighted.

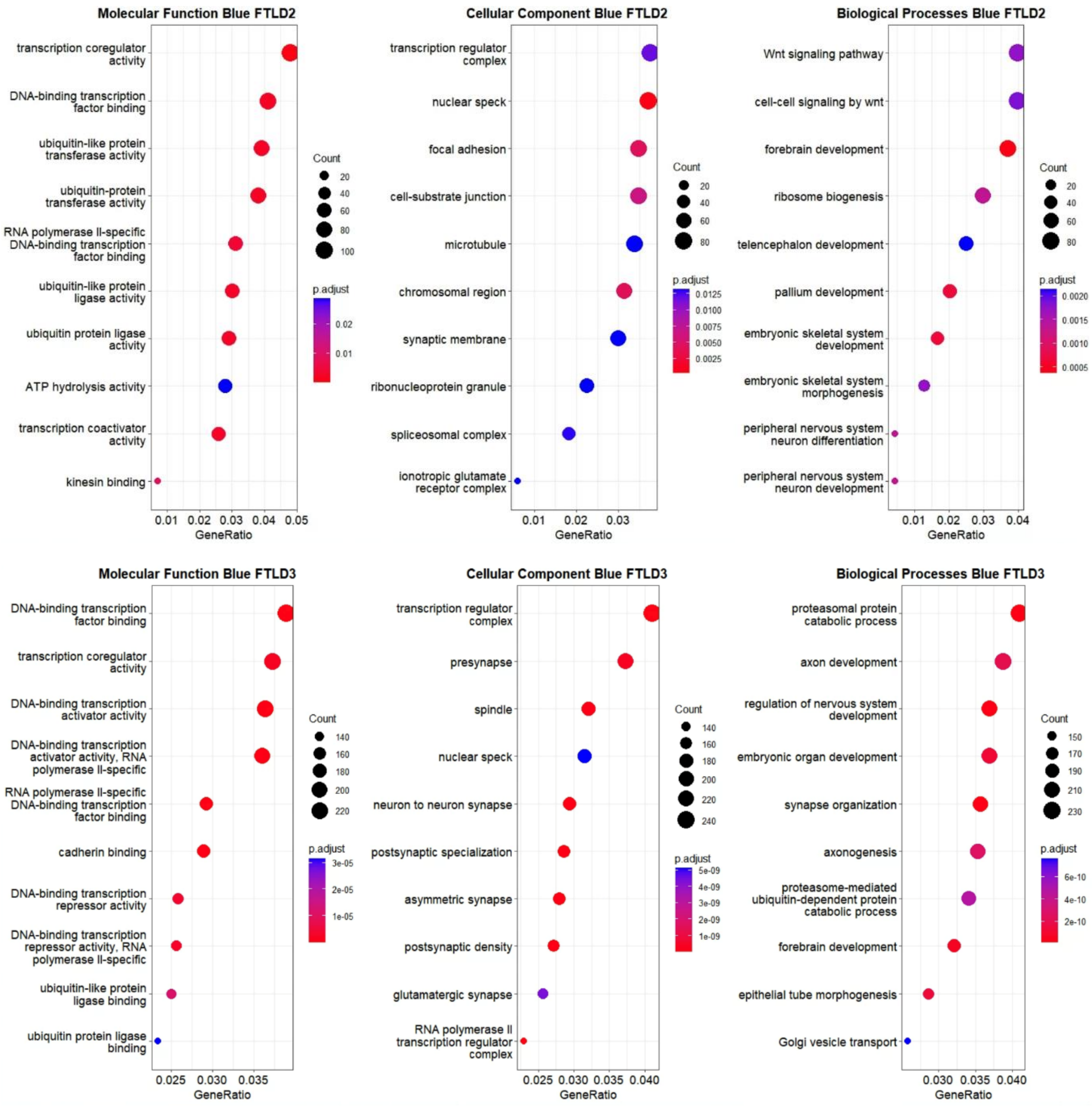


**Supplementary figure 7. Functional enrichment for the co-methylation modules enriched for top meta-analysis hits, FTLD2-blue and FTLD3-blue.** Note: The FTLD1-brown did not retrieve significant enrichment.

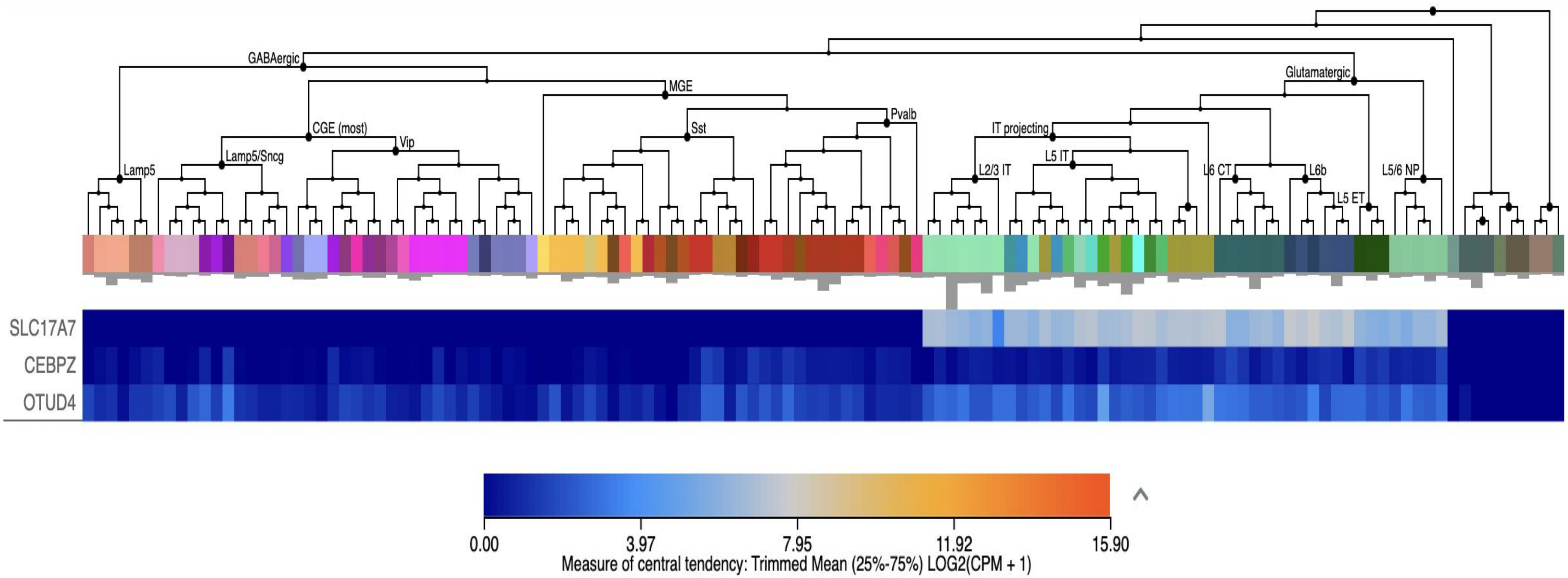


**Supplementary figure 8. Single cell expression patterns of a marker of glutamatergic neurons (*SLC17A7*) and the EWAS meta-analysis hits *OTUD4* and *CEBPZ*.** Single nuclei RNAseq data from the Allen Brain Map (https://celltypes.brain-map.org/)[Bakken et al., 2021[1]]. © 2016 Allen Institute for Brain Science.

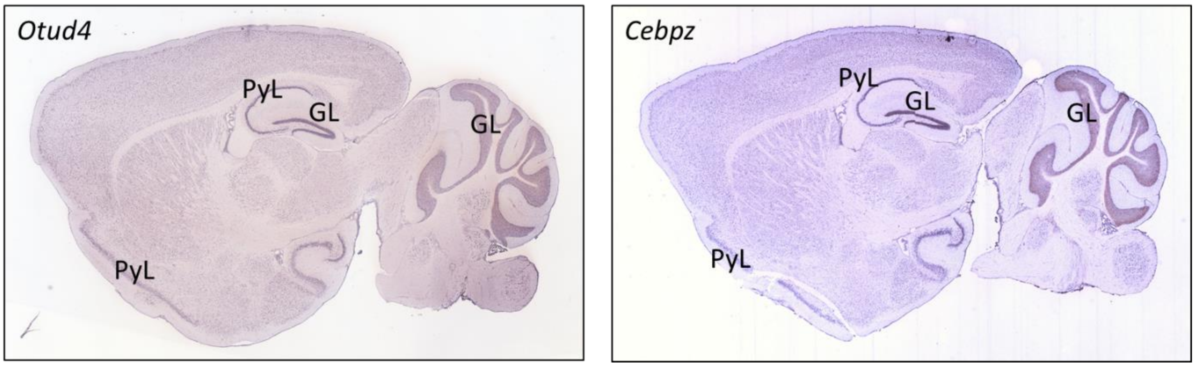


**Supplementary figure 9. Expression patterns of EWAS meta-analysis hits *OTUD4* and *CEBPZ* in mice brains.** Data from the Allen Brain Atlas (http://mouse.brain-map.org) (Lein et al., 2007[3]). PyL – cortical/hippocampal pyramidal layers; GL – cerebellar/hippocampal granule cell layers. © 2016 Allen Institute for Brain Science

**a)**

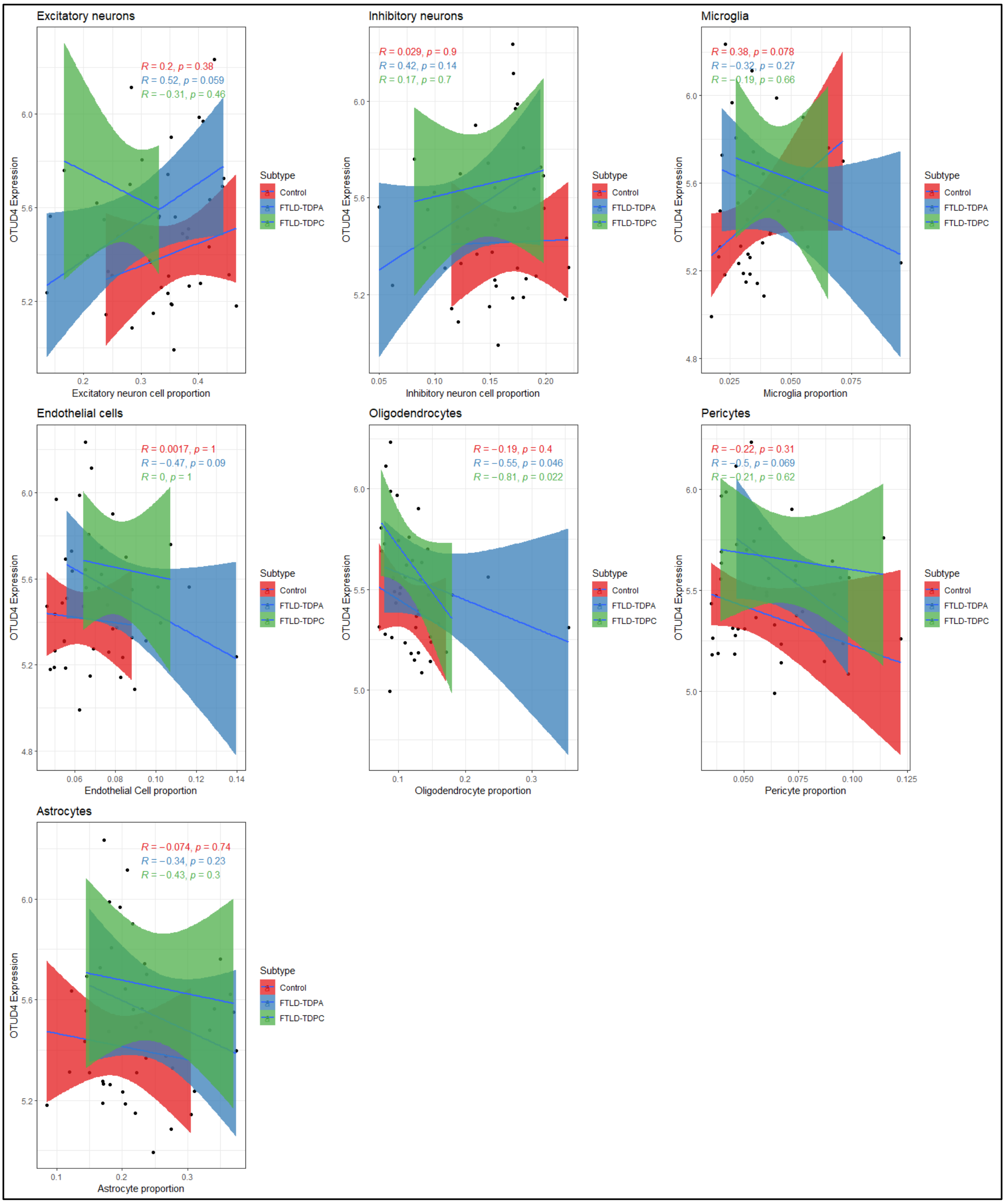
Excitatory neurons
R = 0.2, p = 0.38
R = 0.52, p = 0.059
R = −0.31, p = 0.46
OTUD4 Expression
Excitatory neuron cell proportion
Subtype
Control
FTLD-TDPA
FTLD-TDPC
Inhibitory neurons
R = 0.029, p = 0.9
R = 0.42, p = 0.14
R = 0.17, p = 0.7
OTUD4 Expression
Inhibitory neuron cell proportion
Subtype
Control
FTLD-TDPA
FTLD-TDPC
Microglia
R = 0.38, p = 0.078
R = −0.32, p = 0.27
R = −0.19, p = 0.66
OTUD4 Expression
Microglia proportion
Subtype
Control
FTLD-TDPA
FTLD-TDPC
Endothelial cells
R = 0.0017, p = 1
R = −0.47, p = 0.09
R = 0, p = 1
OTUD4 Expression
Endothelial Cell proportion
Subtype
Control
FTLD-TDPA
FTLD-TDPC
Oligodendrocytes
R = −0.19, p = 0.4
R = −0.55, p = 0.046
R = −0.81, p = 0.022
OTUD4 Expression
Oligodendrocyte proportion
Subtype
Control
FTLD-TDPA
FTLD-TDPC
Pericytes
R = −0.22, p = 0.31
R = −0.5, p = 0.069
R = −0.21, p = 0.62
OTUD4 Expression
Pericyte proportion
Subtype
Control
FTLD-TDPA
FTLD-TDPC
Astrocytes
R = −0.074, p = 0.74
R = −0.34, p = 0.23
R = −0.43, p = 0.3
OTUD4 Expression
Astrocyte proportion
Subtype
Control
FTLD-TDPA
FTLD-TDPC

**b)**

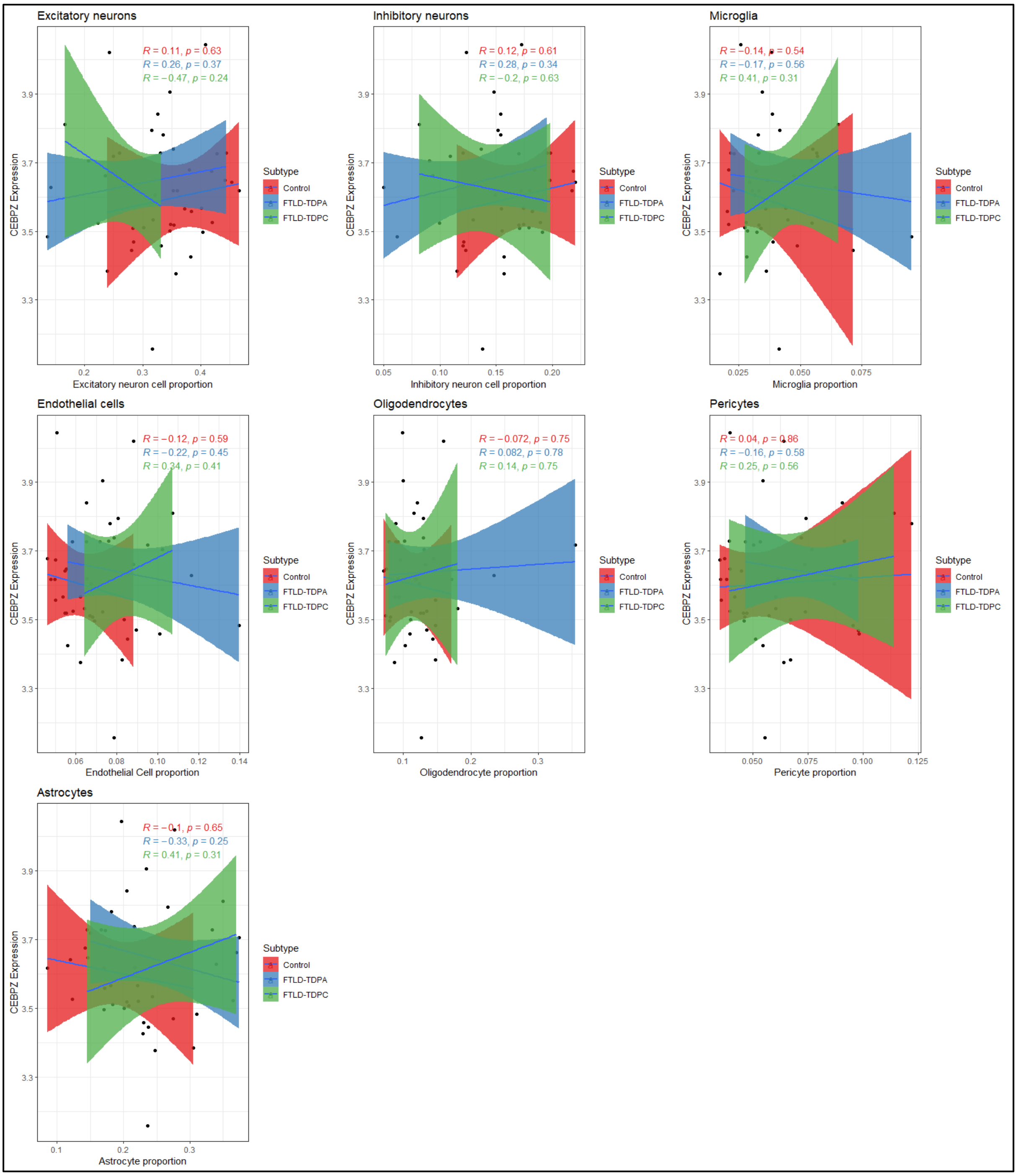


**Supplementary figure 10. Relationship between cellular proportions and gene expression levels of meta-hits. a)** *OTUD4* gene expression vs cell type proportions; **b)** *CEBPZ* gene expression vs cell type proportions. Expression data from Hasan et al. (2022)[3].
